# An Optrode Array for Spatiotemporally Precise Large-Scale Optogenetic Stimulation of Deep Cortical Layers in Non-human Primates

**DOI:** 10.1101/2022.02.09.479779

**Authors:** Andrew M. Clark, Alexander Ingold, Christopher F. Reiche, Donald Cundy, Justin L. Balsor, Frederick Federer, Niall McAlinden, Yunzhou Cheng, John D. Rolston, Loren Rieth, Martin D. Dawson, Keith Mathieson, Steve Blair, Alessandra Angelucci

## Abstract

Optogenetics has transformed studies of neural circuit function, but remains challenging to apply in large brains, such as those of non-human primates (NHPs). A major challenge is delivering intense, spatiotemporally precise, patterned photostimulation across large volumes in deep tissue. Such stimulation is critical, for example, to modulate selectively deep-layer corticocortical feedback projections. To address this unmet need, we have developed the Utah Optrode Array (UOA), a 10×10 glass needle waveguide array fabricated atop a novel opaque optical interposer then bonded to an electrically addressable μLED array. *In vivo* experiments with the UOA demonstrated large-scale, spatiotemporally precise, activation of deep circuits in monkey cortex. Specifically, the UOA permitted both focal (confined to single layers/columns), and widespread (multiple layers/columns) optogenetic activation of deep layer neurons, simply by varying the number of activated μLEDs and/or the irradiance. Thus, the UOA represents a powerful optoelectronic device for targeted manipulation of deep-layer circuits in NHP models.

---

Optogenetics has transformed the study of neural circuit function by allowing for the selective modulation of neural activity on a physiologically relevant timescale^1^. Progress in applying optogenetics to non-genetically tractable models, such as the non-human primate (NHP), has lagged behind that in the mouse^2^. Extending optogenetics to NHP studies is crucial, as, due to their similarity to humans, NHPs represent critical models for understanding neural circuit function and dysfunction^3-6^ and provide an essential technology testbed towards the potential application of optogenetics as therapeutic interventions in humans ^7,8^. The continuing refinement of viral methods for selectively delivering opsins to particular circuits ^9,10^ or cell types^11-13^, is opening up new opportunities to study neural circuits in NHPs^2,14^. Despite these advances, a significant remaining obstacle is the lack of devices for reliably delivering light of sufficient intensity to deep neural tissue across relatively large brain volumes with sufficient spatial resolution to modulate relevant circuit elements.

There are several features of cortical networks that provide both impetus and design requirements for such a device. For example, cortico-cortical feedback connections, which are critical for the contextual modulation of sensory processing^9,15^ and various cognitive phenomena^16,17^, as well as cortico-thalamic feedback projections, arise from deep cortical layers^18,19^. Dissecting these circuits requires selective perturbation of deep layer neurons with high spatiotemporal precision. Moreover, the columnar architecture of the NHP cortex, which extends throughout the cortical layers^20^, requires patterned optogenetic perturbations at the spatial scale of cortical columns through the cortical depth. Methods for high-spatial resolution optogenetics recently developed in smaller animals^21,22^ only allow for stimulation of the superficial layers in the NHP.

Currently, NHP optogenetic experiments mainly follow two light delivery approaches: through-surface illumination and penetrating probes. Surface photostimulation utilizes either a laser- or LED-coupled optical fiber positioned above the cortex^9^, or chronically-implantable surface LED arrays^23^. These approaches enable photoactivation of a large area, but only to a depth of < 1mm, due to light attenuation and scattering in tissue. Furthermore, they result in unintended superficial layer neuron activation and even heating damage at the higher intensities required to reach deep layers^9,24^. In contrast, penetrating optical fibers, integrated with single^25,26^ or multiple^27^ recording probes, allow photoactivation at depths >1mm, but only of a volume a few hundred microns in diameter, and, due to their size and shape, can cause significant superficial layer damage.

To overcome the above limitations, we developed the Utah Optrode Array (UOA), a 10×10 array of glass needle shanks tiling a 4×4 mm^2^ area bonded to an electrically-addressable μLED array independently delivering light through each shank^28,29^. Here, we introduce a second-generation device that incorporates a thin, opaque, optical interposer layer between the needle and μLED arrays that increases light coupling efficiency and virtually eliminates stray light. Furthermore, the entire device underwent a robust encapsulation and testing process to enable *in vivo* testing. Our *in vivo* testing was performed in macaque primary visual cortex (V1), which demonstrated that the UOA allows for spatiotemporally patterned photostimulation of deep cortical layers with sub-millimeter resolution (at the scale of single layers and columns) over a large volume. This selectivity can be scaled up to multiple layers and columns by varying the number of simultaneously activated μLEDs and/or the light irradiance. These results establish the UOA as a powerful tool for studying local and large-scale populations of deep layer neurons in NHP cortex.

## RESULTS

### The UOA: Geometry and Optical Properties

The UOA is based on the geometry of the Utah Electrode Array (UEA)^30^. It is a 10×10 array of penetrating glass optical light guides (needles), with customizable length (up to 2.5mm) and shank width (80-120μm) on a 400μm pitch, tiling 16mm^2^. A custom μLED array fabricated on a GaN on Sapphire wafer is directly integrated with the device, with each electrically addressable 80 × 80μm μLED delivering 450nm light through a single needle (**Fig. 1A-E**). A second 9×9 array of “interstitial” μLEDs is interleaved on the same device for independent surface stimulation (as shown in **Fig. 1B**, but not used in this study). To limit the spatial spread of coupled light, the first generation UOA used a metal pinhole array^28^. Bench testing demonstrated the potential of this device for delivering patterned light at irradiances in excess of activation thresholds across a range of commonly employed depolarizing^31^ and hyperpolarizing^32^ opsins, with a 50% decrease in irradiance within tissue about 200μm from a needle tip^28^, thus providing for depth selectivity. These initial results suggested that direct optogenetic activation through the UOA is on a spatial scale commensurate with the functional architecture of primate cortex.

**Figure 1.**
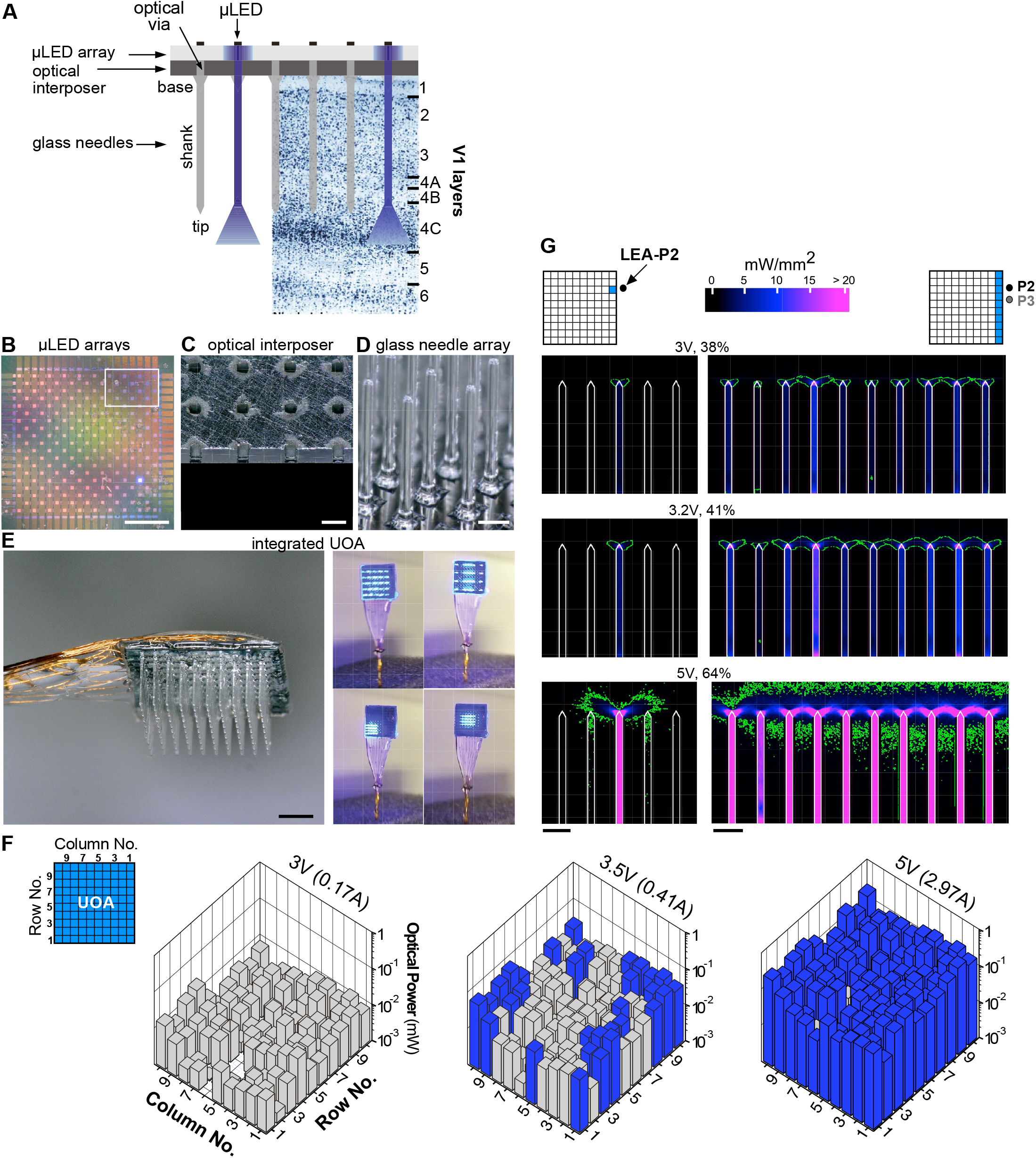
UOA Design and Optical Properties. **(A)** Schematics of UOA design superimposed to a Nissl-stained coronal section of macaque V1 showing cortical layers. The UOA consists of 3 main components: a μLED array (B), an optical interposer (C) and a glass needle array (D). **(B)** Two interleaved μLED arrays on a sapphire substrate are shown in this image; the first 10×10 array is needle-aligned for deep layer stimulation, the second 9×9 interstitial array lies in-between the first for surface stimulation. The interstitial array, although built into the UOA, was not used in this study. Scale bar: 1mm. **(C)** A region of the silicon optical interposer corresponding to approximately the size of the *white box* in (B); the optical “vias” are etched through the silicon and matched to the size of a μLED (80×80μm^2^). Scale bar: 200μm. **(D)** High magnification image of the glass needle shanks bonded to the interposer. Scale bar: 200μm. **(E) <u>Left</u>**: The μLED on sapphire and needle array components are integrated into the final device, wire-bonded, and encapsulated. The image shown is a representative device. The integrated UOA used in this study consisted of 10×10 glass needle shanks, 1.6 mm long (to target deeper layers) and 100-110μm wide, with tip apex angles about 64º. An image of the actual device used in the *in vivo* testing studies, after completion of the experiment and explantation is shown in **Extended Data Fig. 2**. Scale bar: 1mm. **<u>Right</u>**: Example spatial patterns of device operation. **(F)** Average output optical power (in mW) across each needle tip at different drive voltages (currents), when the entire UOA was turned on (*top left inset*). *Blue and gray bars*: needle shanks with estimated tip irradiances above and below, respectively, the 1mW/mm^2^ threshold for ChR2 activation. **(G) <u>Left</u>**: Ray trace model of light spread in cortical tissue when a single μLED (in column 1 and row 8, i.e., the closest to the linear electrode array ––LEA– in penetration 2 –P2– used for the electrophysiological testing experiment, and indicated as a *black dot*) is activated at various input voltages (% of maximum intensity used), with power output calibrated to the bench tests. **<u>Right</u>**: Model of light spread in tissue when all of column 1 (the nearest to the LEA in P2 and P3) is activated at various input voltages. *Green contour* encloses tissue volume within which the light irradiance is above 1mW/mm^2^, the threshold for ChR2 activation. Scale bars: 400μm.

Here we have developed the second generation UOA, which incorporates an optically opaque interposer layer with optical “vias” to eliminate unwanted surface illumination and inter-needle crosstalk (**Fig. 1A, C**; see Online Methods for manufacturing details). This device **(Fig. 1A-E**) was first bench tested (**Fig. 1F**), and from those measurements, *in vivo* optical performance was estimated via ray tracing (**Fig. 1G**). Maps of output power, at each needle tip at different drive voltages, are shown in **Fig. 1F** (**Extended Data Fig. 1** also shows the estimated output irradiances). At 3V, output power and estimated irradiance levels are below the 1mW/mm^2^ threshold for the excitatory opsin *Channelrhodopsin-2* (ChR2) (**Extended Data Fig. 1 and Extended Data Table 1**). Note that defining the irradiance emitted from faceted optrode tips is challenging. For simplicity, in **Extended Data Figure 1B**, we define the irradiance as the emitted optical power divided by the area of the emission surface; however, optical modeling indicates that the emission is non-uniform across the tip surface, with higher irradiance near the tip apex (**Fig. 1G**). There is also variation in emission across the array, due primarily to variations in the resistance (and therefore slope efficiency) of each μLED. At 3.5V, about 30% of the stimulation sites reach or exceed ChR2 threshold (mean optical power ± SD = 0.022 ± 0.013mW; mean irradiance = 0.82 ± 0.49mW/mm^2^), while at 5V, more than 90% of the sites emit above threshold (0.1 ± 0.056mW; 3.79 ± 2.08mW/mm^2^). In principle, software modifications in the matrix driver interface can be made to better equalize stimulation levels across the array.

Using optical ray tracing, we estimated the direct neural stimulation volume (based upon the local irradiance in tissue) as a function of drive voltage and pattern of activated needles to facilitate interpretation of the *in vivo* results (see Online Methods). The left column panels in **Figure 1G** show the stimulation volume, in cross-section, along the first UOA column as produced by the needle (column 1, row 8) nearest one of the electrode penetrations (penetration 2 –P2) in the *in vivo* experiments; the right column panels show the activation volume when all of column 1 is activated. At low drive voltage (∼3V), highly localized stimulation in tissue near the needle tips is produced (as mentioned, the irradiance across the tip surface is non-uniform – concentrated near the apex – explaining why above-ChR2-threshold irradiance levels can be achieved at 3V). At higher voltages (≥ 5V), the stimulation volume overlaps that of adjacent needles, while also extending deeper into tissue. When driving an entire column, at 3V, stimulation localized near each tip is mostly retained, whereas a nearly continuous stimulation volume is obtained at 3.2V due to overlapping intensity patterns. At 5V, the depth of this continuous volume increases, both above and below the tips.

### *In Vivo* Testing: Electrophysiology

We used *in vivo* linear electrode array (LEA) recordings to assess the performance of the UOA for precise modulation of activity in deep layer neurons expressing ChR2. ChR2 and tdTomato (tdT) were expressed in macaque V1 via a mixture of Cre-expressing and Cre-dependent adeno-associated viral vectors (AAV9)^9^. Following a survival period, we recorded multi-unit spiking activity (MUA) using a 24-contact LEA inserted nearby the UOA implanted into a region of dense tdT expression (**Fig. 2A-C; Extended Data Figs. 2-3A**). We performed 3 LEA penetrations (P1-P3), but modulation of neural activity via UOA photostimulation was only detected for P2 and P3 (likely because P1 was farthest from the region of tdT/ChR2 expression; see **Extended Data Fig. 3A**). Below we report data from P2 and P3.

**Figure 2.**
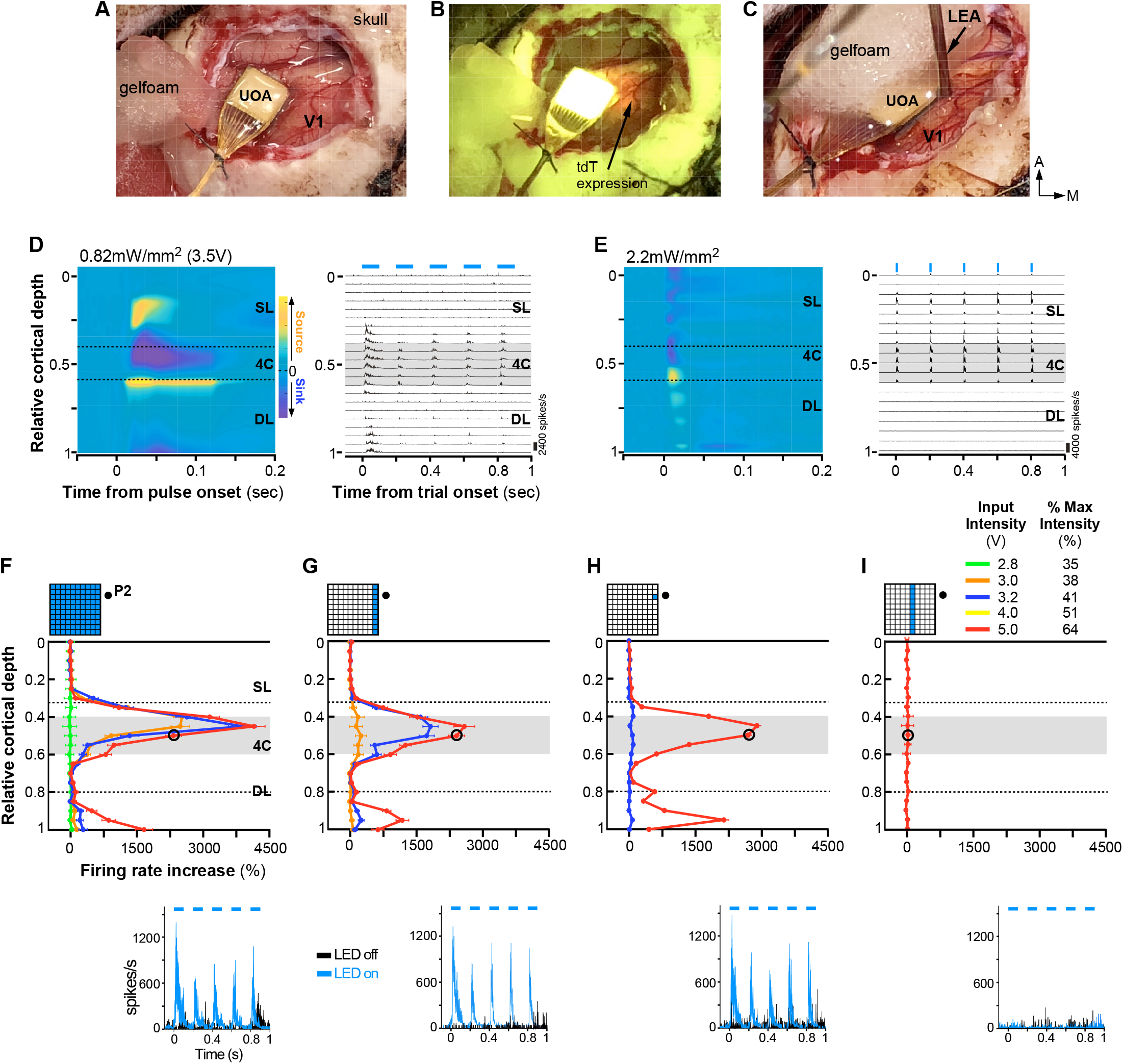
Laminar Distribution of Responses Induced by UOA Photostimulation. **(A)** The UOA used in the *in vivo* experiments inserted in macaque V1. **(B)** Same field of view as in (A) shown under fluorescent illumination to reveal expression of the red fluorescent protein tdTomato (*arrow*). **(C)** Preparation for recording electrophysiological responses to photostimulation. A 24 channel linear electrode array (LEA) was inserted next to the UOA (guide tube protecting array marked “LEA”) slightly angled laterally (towards the UOA) and posteriorly. Here the UOA is partially covered with a piece of Gelfoam. **(D)** Current Source Density analysis (CSD; **Left**) and MUA (**Right**) signals recorded through the depth of V1 in P2 in response to phasic UOA photostimulation (pulse parameters: 100ms pulse duration, 5Hz, 0.82mW/mm^2^; pulse periods denoted as *blue bars* above MUA plot). Here, all 100 needle-aligned μLED sites (“whole μLED array” condition) were activated simultaneously. CSD responses to each 100ms pulse were zero-aligned, while MUA is shown for the full 5Hz pulse train. *The dashed lines in the CSD panel* demarcate the borders of layer 4C (L4C); *the gray shaded region in the MUA panel* delimits the extent of L4C. **(E)** Same as in (D), but following surface photostimulation of V1 via a laser-coupled optical fiber with pulse parameters of 10ms, 5Hz, 2.2mW/mm^2^. **(F-I) <u>Top</u>**: Relative cortical depth of each contact on P2 (*black dot in the insets*) is plotted versus the relative response (% firing rate increase over baseline) to UOA stimulation for different 450nm μLED illumination patterns (*top insets*). Different colored traces are data for different photostimulation intensities (expressed as voltage or percent of max intensity used). *Gray area:* extent of L4C; *dashed lines*: approximate location of the L4A/4B (upper) and L5/L6 (lower) borders. In all panels, error bars represent standard error of the mean. **<u>Bottom</u>**: PSTHs with and without μLED activation are shown for the same contact on the LEA in L4C (marked by the *black circle* in the graphs above) across conditions. *Dashed line in the PSTH*: pulse periods.

### Comparison of Surface and UOA Photostimulation

**Figure 2D** shows neural responses recorded in P2 to simultaneous activation of μLEDs at all UOA sites (whole array condition) at an irradiance level of 0.82 ± 0.49 mW/mm^2^ (pixel-pixel average ± SD) induced by an input intensity of 3.5V (see **Extended Data Table 1**) roughly equivalent to ChR2 activation threshold^31^. To examine the spatiotemporal distribution of responses to UOA stimulation across V1 layers, we first performed a current source density (CSD) analysis of the local field potential (LFP) recorded across the LEA around the time of a UOA pulse (see Online Methods). The CSD reveals the location of current sinks (negative voltage deflections reflecting neuronal depolarization) and sources (positive voltage deflections reflecting return currents) throughout the cortical depth. Current sinks and strong phasic MUA in response to UOA stimulation were mostly localized to layer (L) 4C and the lower part of the deep layers, with L4C activity preceding activity in deeper layers (**Fig. 2D**). This suggests that the UOA needle tips closest to P2 terminated in L4C and that, at these low photostimulation intensities, light spread remained close to the UOA tips. In contrast, at the highest intensity, light spreads farther into deeper layers (**Extended Data Fig. 4A-B**). Importantly, this qualitatively distinct laminar pattern of neural activation could not be explained by thermal artifacts (**Extended Data Figs. 5-6**). Additional analysis demonstrated that response onset latency and onset reliability were lowest and highest, respectively, for the P2 contacts located in L4C. Combined with postmortem histological assessment, this confirms the UOA needle tips closest to P2 were located in L4C (**Extended Data Fig. 3A, B Right**). Comparison of these laminar activity patterns elicited by UOA photostimulation with that elicited by direct surface photostimulation in a different animal at a slightly higher irradiance (2.2mW/mm^2^) revealed a sharp dissociation. Specifically, surface stimulation of ChR2 evoked responses starting in superficial layers and terminating in L4C **(Fig. 2E**).

### UOA Stimulation Parameters Can Be Tuned to Achieve Laminar Specificity

To assess the impact of UOA stimulation on MUA we varied: (i) the spatial pattern of UOA stimulation (single μLED sites, entire columns, or the entire device) and (ii) stimulation intensity across these spatial patterns. In all conditions, we used phasic stimulation (5Hz, 100 msec pulses for 1 sec with 1.5-21 sec inter-trial intervals, with the longer intervals used at the higher stimulation intensities) with a slow on/off ramping to eliminate the potential of any electrical artifacts induced by capacitive coupling at the array/tissue interface^33^. As an example, **Figure 2F-I** shows responses from P2. As indicated by an analysis of firing rate increase across layers induced by activating a single μLED at different sites along column 1, the UOA needles closest to P2 were those in rows 8 and 9 (C1-R8, C1-R9), and their tips terminated into L4C (**Extended Data Fig. 3B Left**). The laminar distribution of MUA in P2 varied in amplitude across conditions, but was reliably confined to deeper layers. By varying the spatial pattern of stimulation and/or the stimulation intensity, MUA could be confined to single layers or spread across multiple layers. For example, activation of the whole UOA (**Fig. 2F**) at intensities > 2.8V and up to 5V evoked a MUA peak within L4C (where the needle tips nearest to P2 terminated). This peak increased in magnitude with increasing stimulation intensity. Increasing this intensity range further (4-5V) led to a second, smaller, MUA peak in L6 (but not L5). In macaque V1, L4C projects to both L5 and L6^34^, but its net effect is to suppress the former^35^ and activate the latter^36^, consistent with the interpretation that at the higher light intensities, lack of L5 responses and increases in L6 responses may have resulted from synaptic spread from optogenetically-activated L4C neurons. Below, we provide evidence supporting this interpretation. At even higher intensities, neural activity increased in L4C through L6 likely via direct activation of the deeper layers due to light scattering through a larger volume (**Extended Data Fig. 4C Left**). Although thermal artifacts could not explain the findings at the highest intensity tested with our stimulation parameters (**Extended Data Figs. 5B**,**6**), in general lower stimulation intensities should be favored in experiments, particularly when the entire UOA is activated and shorter intertrial intervals are used. This is because heat-induced perturbations in firing rates can occur at higher intensities during the inter-trial period (**Extended Data Fig. 5A**) and potentially affect trial-specific responses at shorter inter-trial intervals than those used in our study (as suggested in **Extended Data Fig. 6**).

Activation at 5V evoked similar laminar patterns and magnitudes of MUA irrespective of whether a single μLED, an entire column nearest the LEA, or the whole UOA were illuminated (**Fig. 2F-H**). However, at lower photostimulation intensities, firing rate increased with the number of activated μLEDs (e.g., compare blue curves in **Fig. 2F-H**), and higher intensities (>3.2V) were required to modulate neural activity via a single μLED (**Fig. 2H**). Moving the μLED activation column a distance of 1.6mm on the UOA (from column 1 to 5) resulted in a 10-fold reduction in MUA amplitude (**Fig. 2I**), and increases in firing rates in L4C were observed only at the highest intensities used (7.8V; **Extended data Fig. 4C Right)**. No increase in firing rate could be evoked by activation of an entire column beyond this distance or of a single μLED in column 1 beyond a similar distance on the UOA (row 4; corresponding to a distance from the LEA of 2.6-2.7mm estimated from postmortem histology) even at the highest intensity used (7.8V, **Extended Data Table 1**).

### Tangential Extent of Responses Induced by Photostimulation via the UOA

An analysis similar to that performed for P2 allowed us to determine the location of P3 relative to the UOA, and to establish that μLED C1-R7 was the closest to P3 and its tip terminated in the superficial layers (**Extended Data Fig. 3C)**.

We next investigated whether the MUA across LEA contacts was tuned to the spatial site of UOA stimulation. To estimate MUA selectivity for stimulation at UOA sites between columns 1-5 and rows 3-10, we fit a multiple linear regression model to the MUA recorded at each LEA contact, with row, column, and intensity (V) as independent variables (see Online Methods). We included in this analysis only contacts on which there was a significant difference in firing rates during the stimulation and control periods for at least one of the row or column conditions (ANOVA, all p < 0.01). On average, including a quadratic term explained more of the variance in the MUA response (mean R^2^ ± SD: 0.58 ± 0.14 vs. 0.31 ± 0.11 for a linear model; Kolmogorov-Smirnov, p < 10^−7^). **Figure 3A, E** shows plots of fitted MUA for 3.5V single-μLED photostimulation for the contact in P2 and P3 that showed the greatest relative response modulation. We normalized each contact’s fitted responses to the peak, and averaged across contacts to determine whether MUA preferred stimulation at different UOA sites on different LEA penetrations (**Fig. 3B, F**). Consistent with our prior assessment (**Extended Data Fig. 3A-C**), the peaks for P2 contacts tended to cluster mostly near C1-2/R8-9, while those for P3 contacts clustered mostly near C1-3/R4-7. The spatial pattern of peak activity across the LEA suggested that, particularly for P3, the LEA was inserted at a slightly oblique angle. Peak locations differed significantly across the two penetrations (ANOVA, p<0.01).

**Figure 3.**
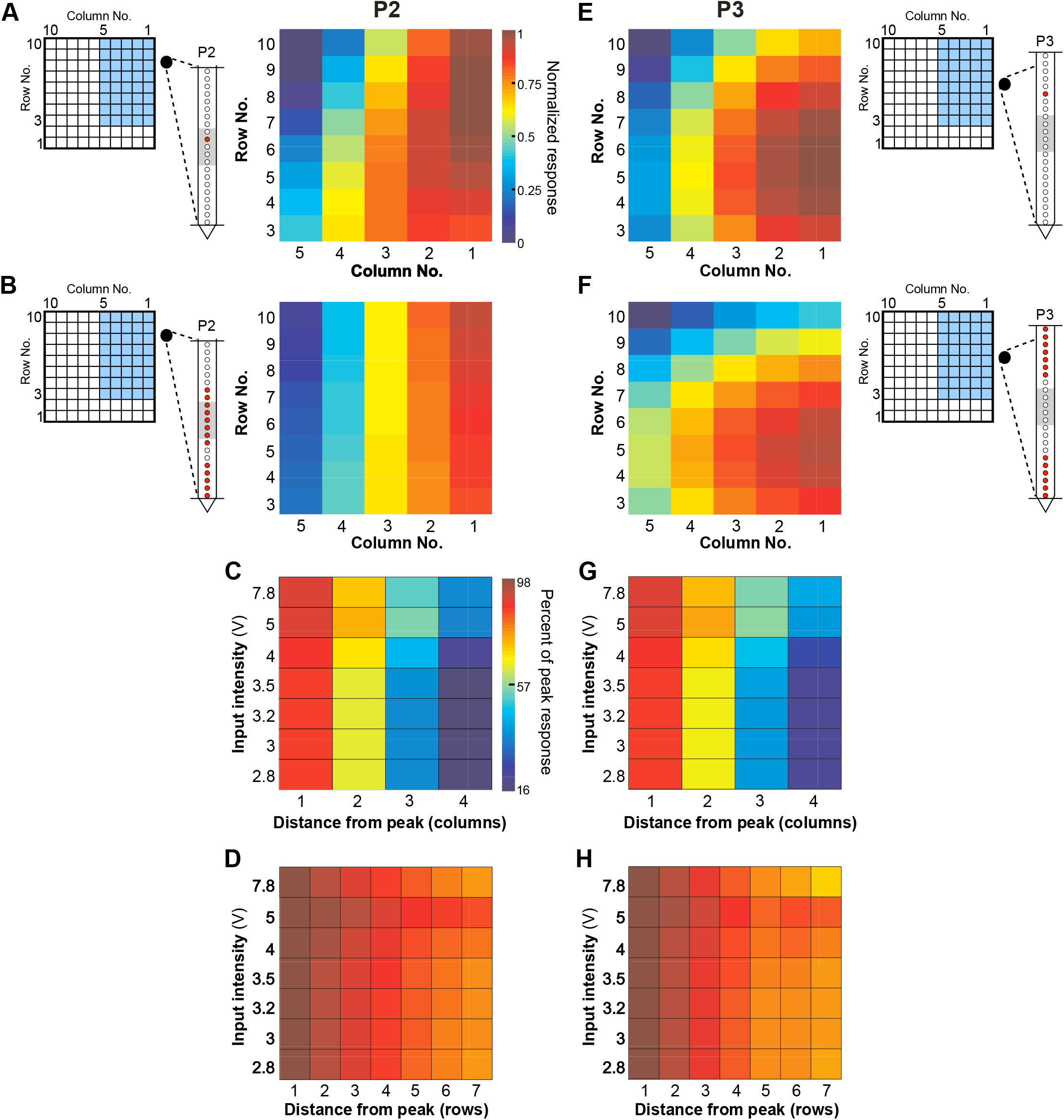
Tangential Extent of Responses Induced by UOA Photostimulation. **A)** Examples of model fits to single μLED photostimulation for the contact from P2 showing the largest relative response increase across these stimulation conditions. This contact preferred stimulation in the proximal UOA columns 1-2, at sites closer to the top of the device (rows 9-7). The schematics on the left of the UOA and of the LEA-P2 indicates as *blue shading* the UOA sites represented in the heat map, and as a *red dot* the contact on the LEA whose response is mapped on the right. The *horizontal lines* and *gray shading* on the LEA schematics mark the pial and white matter, and L4C boundaries, respectively. Color scale applies to panels (A-B, E-F).(B)Average normalized fitted responses across all responsive contacts in P2 (*red dots* in schematics of LEA to the left). **(C)** Change in response in the column direction for P2. Average relative response amplitude (% of peak model-fitted response) is plotted as a function of stimulation intensity and distance along a straight line extending from the preferred UOA site in the column direction, and sorted by input intensity. Data averaged across all responsive contacts.**(D)** Change in response in the row direction for P2. Average relative response amplitude (% of peak response) is plotted as a function of stimulation intensity and distance along a straight line extending from the preferred UOA site in the row direction. Data averaged across all contacts. **(E-H**) Same as in (A-D) but for P3.

The data in **Figures 2 (panels G, I)** and **Figure3 (panels A, B, E, F)** indicated MUA amplitude decreased with increasing distance between photostimulation and recording sites. To quantify this observation, and better characterize the extent of photostimulation-evoked responses across the tangential domain of cortex, we examined MUA amplitude as a function of distance on the UOA (in a straight line extending along either the row or the column axis) from the site that evoked the peak response (**Fig. 3C-H**). As is evident from the steeper decrease in responses along the column versus the row axis, as well as the difference in relative response across stimulus intensities, there was a significant effect of UOA axis and input intensity on relative response (ANOVA, both p < 10^−21^), as well as a significant difference across penetrations (ANOVA, p < 10^−14^). Finally, there was a significant interaction between intensity and UOA axis as well as UOA axis and penetration (ANOVA, both p < 0.01). These results indicate that the response decrease from peak is greater in the column versus the row direction, that intensity has a different effect on this drop-off in the row versus column directions, and that this differed across penetrations. For example, in the column direction, at 2.8V intensity MUA dropped to 16% of peak at a distance of 1.6 mm from peak, but at ≥ 5V it dropped to 50% at the same distance (**Fig. 3C-G**). Instead, in the row direction, at 2.8V MUA dropped to 80% of peak at a distance of 2.8mm, and to 90% at ≥ 5V (**Fig. 3D, H**). The difference in response drop-off with distance in the column vs. row directions is likely explained by the greater differences in irradiance, for a given input intensity, along the column as compared to the row axis (see **Extended Data Fig. 1**).

In summary, the spatial spread of MUA along the tangential domain of cortex varied according to UOA stimulation site and intensity. Importantly, the extent of this spread was more limited at lower intensities, suggesting that increasing intensity increased the volume over which cells were optogenetically activated, consistent with the model simulations in **Fig. 1G**.

### UOA Activation Parameters Can Be Tuned to Activate Distinct Cortical Networks

Given the spatial separation between the LEA and the UOA (∼1-1.1mm for P2 and 700-800μm for P3, based on histology; **Extended Data Fig. 3A**), the reported sharp falloff in light intensity over short distances in tissue^37,38^, and our bench estimates of light spread from the UOA tips^28^ (see also **Fig. 1G**), we reasoned that the evoked MUA we recorded was likely relayed to the recorded neurons indirectly, via activation of ChR2-expressing cells nearby UOA needle tips. To examine this possibility, we measured the onset latency of evoked MUA across layers.

Example latency data from P2 are shown in **Figure 4A**. Here, the UOA stimulus was a single μLED (C1/R8/5V) nearest the recording location. The fastest evoked response occurred in mid layers with an onset latency (see Online Methods) of about 15ms. Deep layer response onset (mean ± s.e.m: 30 ± 7ms) lagged that in mid-layers, as would be expected if optogenetic activation first propagated through L4C before being synaptically relayed to deeper layers, via L4C-to-L5/6 connections. Averaged PSTHs for the peri-pulse period on one example L4C and one L6 contact are shown in **Figure 4B**. There was a significant pulse-by-pulse difference in onset latency across contacts (ANOVA, p < 10^−30),^ as well as a significant pairwise difference across these two LEA recording sites (Tukey HSD test, p < 10^−6^; **Fig. 4B Right**).

**Figure 4.**
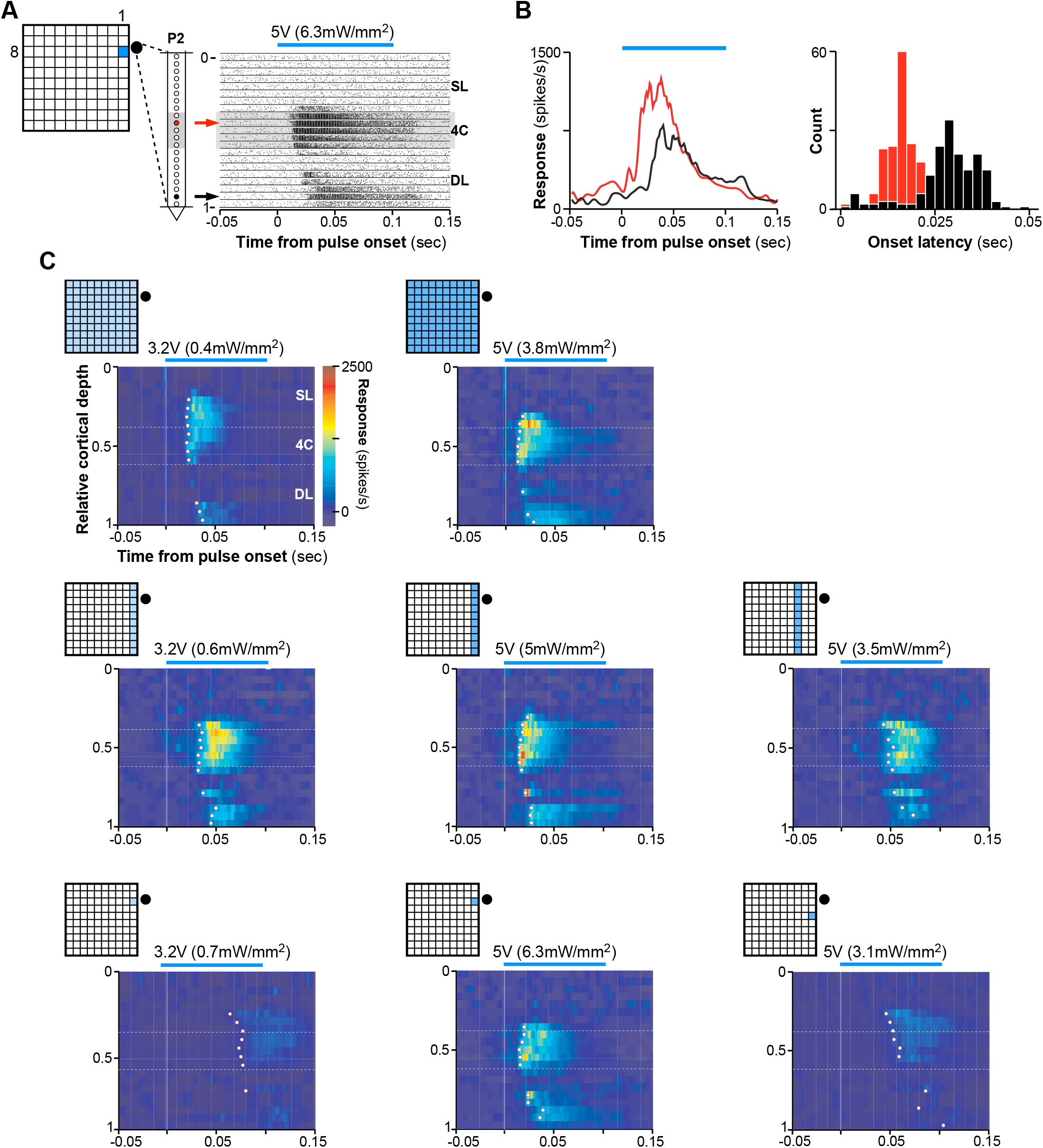
Onset Latencies Reveal Local Networks Activated by Focal Optogenetic Modulation. **(A) <u>Left</u>**: Schematics of UOA stimulation through a single μLED site (C1-R8) and of relative LEA position in P2. **<u>Right</u>**: Pulse-aligned raster plots for all 21 channels on the LEA through the depth of V1. *Black lines* separate data from different channels. *Gray shaded region:* channels in L4C. *Blue line above plot*: 100ms pulse period at the input voltage (irradiance) indicated. *Red and black arrows denote example contacts in L4C and 6, respectively*. A graded shift in MUA onset latency is apparent. **(B) <u>Left</u>**: Pulse-aligned PSTHs for the two channels indicated by arrows in the raster plot in (A). Responses are plotted as baseline-subtracted firing rate versus time. Response onset latency at the L6 contact (35ms) clearly lagged that on the L4C contact nearest the UOA needle tips (17ms). **<u>Right</u>**: Histograms of pulse-by-pulse onset latencies for the two example contacts. **(C)** Heatmaps of MUA (firing rate) through the depth of V1 during the peri-pulse period, for the UOA stimulation condition indicated by the insets at the top left of each plot. Stimulation intensity (average irradiance) is reported above each plot. The firing rate color scale applies to all panels. *White dots* mark the onset latency (estimated from the mean PSTH- see Online Methods) for each contact that was significantly responsive to UOA stimulation.

**Figure 4C** shows average peri-pulse PSTHs across all LEA contacts as a function of normalized cortical depth for exemplary whole array (top panels), single column (middle panels), and single μLED (bottom panels) stimulation at different intensities or μLED-LEA distances. Increasing total stimulus area at lower intensities (panels in the left column of **Fig. 4C**) increased the number of responsive contacts and the amplitude of driven responses, and shortened onset latencies. At higher intensities (5V, middle column), there was little change in these measures across large differences in total stimulated area. Decreasing the stimulus intensity for a fixed area (middle to left columns in **Fig. 4C**), or increasing the separation between the stimulated UOA site/s and the LEA for a fixed stimulus intensity (middle to right panels in the center and bottom rows of **Fig. 4C**) increased onset latencies across all contacts (mean latency ± s.e.m at 5V and 3.2V: 17 ± 1.7ms and 25.4 ± 2ms, respectively, whole array condition; 19.8 ± 1.4ms and 37.5 ± 1.9ms, C1 condition; 21.4 ± 2.3ms and 74.1 ± 1.6ms, C1-R8 condition; mean latency ± s.e.m at 5V: 47.6 ± 4.3ms and 59.4 ± 4.1ms for C3 and C1-R6 conditions, respectively). Calculating onset latency on a pulse-by-pulse basis and looking at the effects on latency of cortical depth, stimulation pattern, and stimulation intensity, we observed significant main effects of pattern and intensity, as well as significant two-way and three-way interactions between all three factors (ANOVA, all p < 10^−4^). Limiting our analysis to each pattern, we observed a significant effect on latency of intensity and distance from the LEA for the single column conditions in **Fig. 4C** (ANOVA, all p < 10^−4^), and a significant effect of distance for the single μLED conditions (ANOVA, p = 0.03). Furthermore, in many conditions, pairwise comparisons across contacts revealed a significantly delayed response onset in deep layers relative to mid-layers for most conditions in **Figure 4C** at 5V, and for some conditions at 3.2V (Tukey HSD, all p < 0.01; **Extended Data Fig. 7**); this time-lag varied with intensity and separation between stimulation and recording sites, increasing at lower intensities and greater distances. There was also a significant difference in onset latency between mid- and superficial layers in some conditions (C1 at 5V, whole array at 5V and 3.2V; Tukey HSD, all p < 0.01; **Extended Data Fig. 7**). Notably, however, when the whole μLED array was stimulated at the highest intensity (7.8V), there was no significant difference in onset latencies between deep and middle layers, again suggesting the former were directly activated by light spreading through deeper tissue (**Extended Data Figs. 4D and 7**).

To quantify these effects across the population (n= 33 significantly responsive contacts, across 2 LEA penetrations), we first calculated the distance between each LEA contact and the contact with the shortest onset latency, and plotted this distance versus onset latency, separately for each unique combination of UOA stimulation site(s) and intensity. Similar to the P2 data shown in **Fig. 4C**, the population data showed 2 main effects. (1) Onset latency decreased significantly across all contacts with increasing stimulation intensity (ANOVA, main effect of intensity, all p<0.01; **Fig. 5A, 5B Left, 5C Left**) and proximity to the recording LEA site (ANOVA, main effect of row or column on UOA, all p<10^−4^; right panels in **Fig. 5B and 5C**). (2) Onset latency increased significantly with contact distance on the LEA from the fastest contact (**Fig. 5A-C**, main effect of distance on the LEA, ANOVA all p<0.01), suggesting that the more distant contacts were activated indirectly via interlaminar networks. However, for stimulation of the whole UOA at higher intensity (7.8V), evoked responses had similar onset latencies across the LEA (thus, across V1 layers; **Extended Data Figs. 4E, 7** top right).

**Figure 5.**
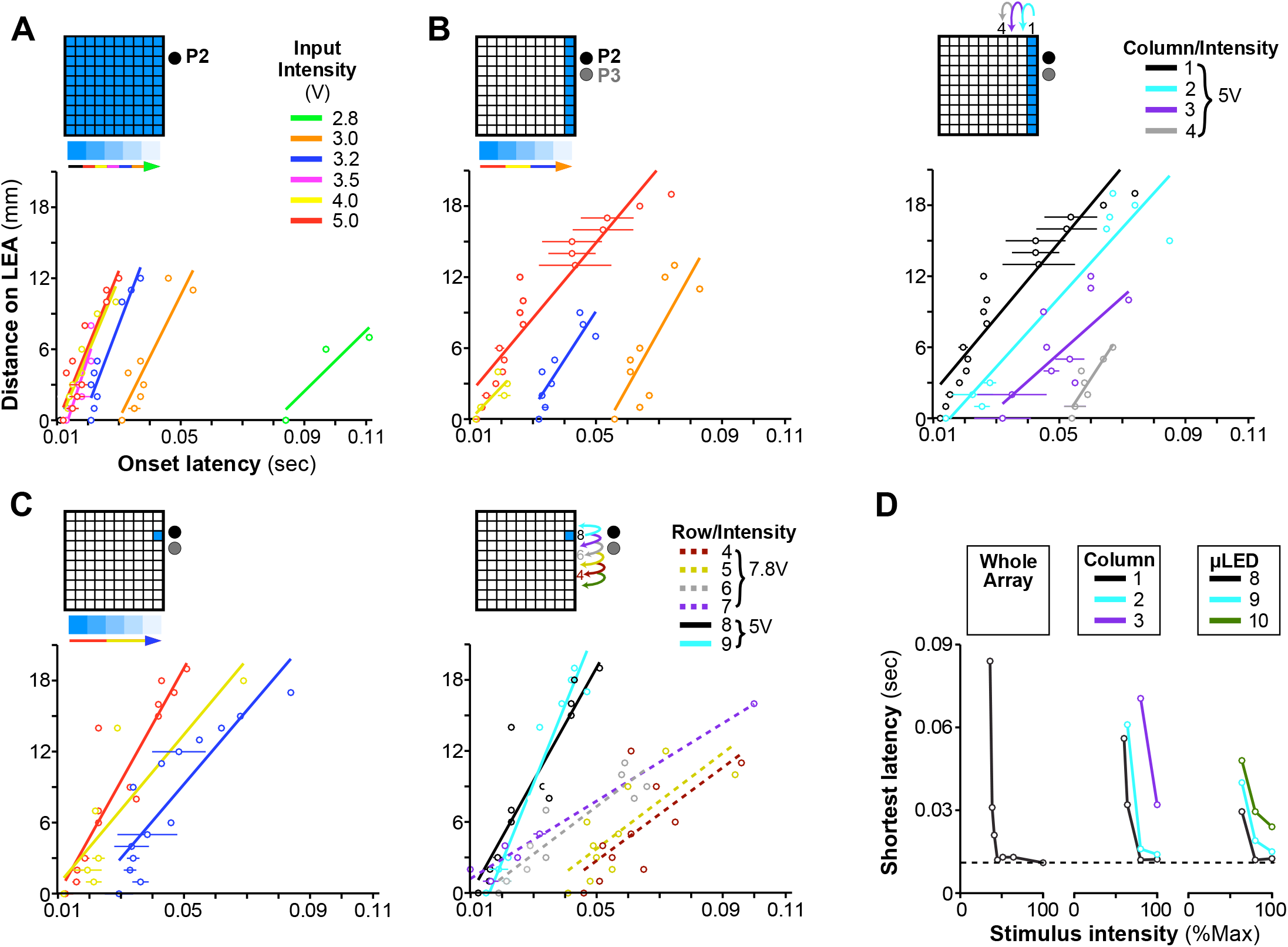
Population Onset Latencies as a Function of UOA Stimulation Intensity and Spatial Pattern. **(A)** Distance on the LEA of each contact from the contact with the fastest onset latency is plotted against onset latency; lines are linear fits. Each line is from simultaneous stimulation throughout the whole μLED array at each indicated intensity. **(B) <u>Left</u>**: Effect of varying photostimulation intensity for a fixed column (C1). **<u>Right</u>**: Effect of varying stimulated column (C1 to C4) for a fixed photostimulation intensity (5V). Either lowering intensity for a given column or increasing the distance between an activated column and the LEA had similar effects on the latency of network activation. **(C)** As in (B), but for a single μLED stimulation condition. **<u>Left</u>**:, photostimulation intensity was varied for a fixed μLED (C1-R8); **<u>Right</u>**: the stimulated μLED was varied along column 1 (from row 3 to 9) at a fixed intensity (5V for μLEDS in rows 8-10, but 7.8V for those in rows 4-7, as lower intensities did not evoke a response from many of these latter μLEDs). **(D)** The shortest onset latency across all intensities (here expressed as percent of max-see legend in **Extended Data Fig. 4C** for corresponding input voltage) is plotted for the whole array condition (**Left**), and selected columns (**Middle**) or μLEDs (**Right**). In all panels, error bars represent standard error of the mean.

Across the three categories of UOA stimulation (whole array, column, and single μLED), only for the whole array and single μLED conditions did we observe a significant interaction between the effects of distance along the LEA and UOA photostimulation intensity on onset latency (**Fig. 5A, 5C Left**; both p < 0.05, ANOVA). In these conditions, lowering photostimulation intensity decreased the slope of the curves, indicating that the difference in onset latency with distance on the LEA increased at lower intensity. Additionally, for the single μLED condition, we also observed a significant decrease in the slope of the curves when stimulating at increasing UOA-LEA separation, but only when we moved the single μLED stimulus to sites that were far enough from the LEA to necessitate stimulation at the very highest powers used to elicit any response (dashed lines in **Fig. 5C Right**, μLED in rows 4-7; ANOVA, LEA distance × UOA row × intensity interaction, p < 10^−3^). For the single column condition, there was no significant interaction between contact distance and either photostimulation intensity or UOA-LEA separation (**Fig. 5B**; ANOVA, all p > 0.09). Importantly, across all three photostimulation patterns (whole array, single columns, and single μLEDs) there was remarkable similarity in the timing of the fastest responses (**Fig. 5D**). Both increasing stimulus area and stimulating at UOA sites closer to the recording locations reduced the light intensity necessary to evoke responses at this latency, but did not result in shorter latencies. This is further evidence that the evoked MUA nearby LEA contacts was relayed indirectly following optogenetic activation at UOA tips, and that the timing of this activation depended upon both the location and area of optogenetically activated inputs.

In summary, by varying photostimulation intensity and/or number of stimulated sites, the UOA allows activation of single or multiple layers, while by varying the spatial separation between the site of UOA stimulation and that of the recording, the UOA allows investigations of local vs long-range intra and interlaminar circuits.

### *In Vivo* Testing: c-Fos Expression

To validate the performance of the UOA for large-scale photostimulation, we measured changes in c-fos expression, an immediate early gene whose expression rapidly increases when neurons are stressed or activated^39,40^. C-fos protein expression can be used as an indirect measure of the spatial pattern of neural activation. We analyzed patterns of c-fos expression using immunohistochemistry (IHC) (see Online Methods) in two control and two experimental hemispheres from 3 animals.

In one experimental case (MK414-RH), a “passive” UOA (lacking an integrated μLED array) was implanted in a ChR2/tdT-expressing region of V1 (**Fig. 6A-B**). We stimulated the deep layers through a subset of needles, using a collimated, fiber-coupled, 473nm laser, while shielding from light surrounding cortex and portions of the UOA (see Online Methods). Histological analysis revealed that the UOA in this case was inserted at an angle (due to brain curvature), its needle tips ending at the bottom of the superficial layers, anteriorly, and in progressively deeper layers, posteriorly (most tips being in L4C, only the most ventral ones reaching L6) (**Fig. 6A-B**). C-fos positive (c-fos+) cells were found throughout V1 (**Fig. 6A, C, D**), as well as in V1 recipient extrastriate areas, including V2 (**Fig. 6A, C, D**), V3, and MT (not shown)). This extensive pattern of elevated c-fos expression was likely induced by direct optogenetic activation and indirectly via synaptic activation. To test this hypothesis, we repeated the experiment in a different animal (MK422-RH) in which we greatly reduced glutamatergic neurotransmission via application of the AMPA receptor antagonist NBQX to ChR2-expressing cortex prior to passive-UOA insertion and photostimulation. Most of the UOA’s needle tips, in this case, only reached the bottom of the superficial layers (**Fig. 6E-F**). We also performed two additional experiments, to control for the potential of elevated c-fos expression being induced by either UOA insertion or stray photostimulation, respectively. In case MK414-LH, we inserted a passive UOA in the supplementary motor area (SMA) not expressing ChR2, and euthanized the animal 4 hours later without stimulating through the array. Histological analysis revealed that the UOA was fully inserted in this case (tips reaching L5; **Fig. 6I**). In case MK421-RH, instead, we only performed surface photostimulation of SMA cortex not expressing ChR2 and no UOA insertion (**Fig. 6K**).

**Figure 6.**
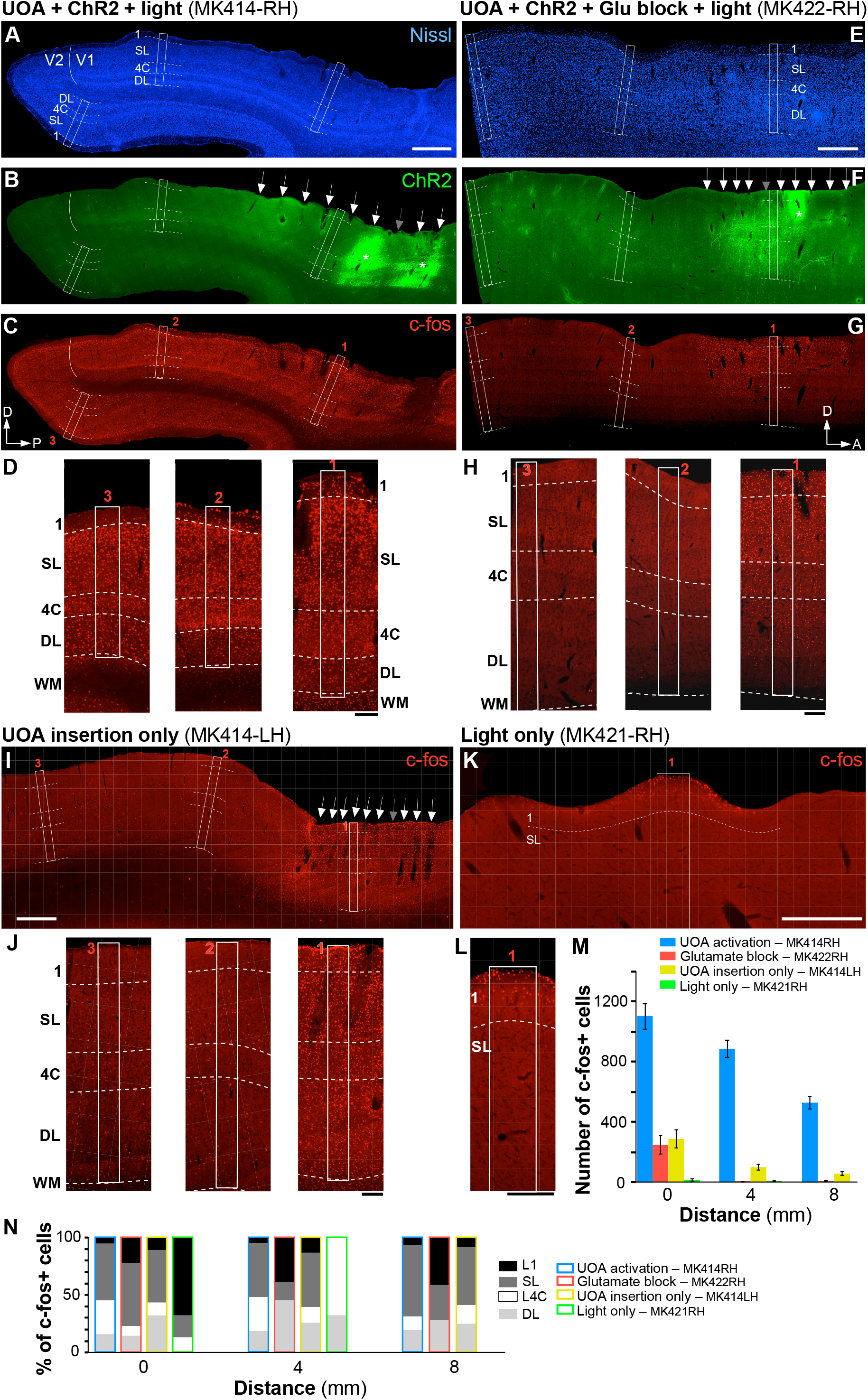
Local Optogenetic Activation Through the UOA Spreads Through Cortico-Cortical Networks. **(A-C)** Case MK414-RH (UOA activation). The same sagittal section encompassing parts of V1 and V2 is shown under 3 different fluorescent illuminations, to reveal Nissl stain (A), tdT/ChR2 expression (B; the red tdT fluorescence was converted to green for purpose of illustration), and c-fos IHC (C). *White solid contour*: V1/V2 border; *dashed contours:* layer boundaries (layers are indicated); *white boxes:* ROIs (numbered 1-3 in panel C) where c-fos+ cells were counted. *White Arrows in (B)* point to the visible damage caused by each UOA needle, while the *gray arrow* points to the likely location of one of the UOA needles that did not cause visible damage in this section. *Asterisks in (B)* mark the core of the viral injections, and sites of highest tdT/ChR2 expression. *P:* posterior; *D:* dorsal. C-fos expression in this case is observed throughout all layers (local) and across cortical areas (long-range). Scale bar in (A): 1mm (valid for A-C). **(D)** Higher magnification of c-fos IHC in and around each ROI. Scale bar: 0.2mm. **(E-H)** Case MK422-RH (Glutamate block). Same as in (A-D) but for a different case in which an AMPA receptor antagonist was injected into the SMA prior to UOA insertion and photostimulation. The sagittal section is from the SMA. *D:* dorsal; *A:* anterior. Scale bars: 1mm (E, valid for E-G); 0.2mmm (H). Blocking AMPA receptors demonstrates that initial optogenetic activation is limited to the stimulated layers in the region of UOA insertion. **(I-J)** Case MK414-LH (UOA insertion-only). C-fos IHC in a sagittal section of SMA cortex (I) and at higher magnification in and around each ROI used for cell counts (J), in a case which only received UOA insertion in cortex not expressing ChR2, and no photostimulation. Scale bars: 1mm (I), 0.2mm (J). **(K-L)** Case MK421-RH (Light-only). Same as in (I-J), but for a control case in which SMA cortex not expressing ChR2 only received surface photostimulation via an optical fiber-coupled laser and no UOA insertion. Here only one ROI is shown at higher magnification to reveal the few labeled cells in L1. Scale bars: 0.5mm (K), 0.2mm (L). The increases in cFos expression seen after full UOA activation of ChR2-expressing cortex cannot quantitatively be explained by device insertion or surface illumination. **(M)** Average number of c-fos+ cells across sections used for quantification, as a function of distance from the center of UOA insertion for the 4 different cases. Error bars: s.e.m. **(N)** Distribution of c-fos+ cells across layers at each distance.

To quantify c-fos expression across our various manipulations, we counted c-fos+ cells in 3 regions of interest (ROIs) encompassing all cortical layers, one centered in the region of UOA insertion and/or light stimulation, the other two located 4 and 8 mm, respectively, from the first (*white boxes* numbered 1-3 in **Fig. 6A-L**; see Online Methods). **Figure 6M** plots the average number of c-fos+ cells across samples, as a function of distance from the UOA insertion site, while **Figure 6N** shows the laminar distributions of c-fos+ neurons at each distance. We found significant local (involving all layers) and long-range c-fos expression only when photostimulation of ChR2-expressing cortex was performed via the UOA (MK414-RH; Fig. **6C-D, M-N**). Blocking glutamate neurotransmission prior to photostimulation prevented long-range c-fos expression, and reduced its expression by 5 fold in the area of UOA stimulation, where it was largely confined to the directly stimulated layers (mostly superficial) near the UOA tips (MK422-RH; **Fig. 6G-H, M-N**). UOA insertion-only led to as much local c-fos expression as the glutamate block case, but to greater interlaminar (involving all layers), as well as intra- and inter-areal long-range spread (MK414-LH; **Fig. 6J, M-N**), suggesting that neurons activated by the insertion trauma also indirectly activated downstream networks. Finally, surface photostimulation of cortex not-expressing ChR2, without UAO insertion, caused virtually no c-fos expression, except for a few cells in L1 and upper L2 (MK421-RH; **Fig. 6L-N**). Statistical analysis (one way ANOVA with Bonferroni corrected post hoc comparisons) revealed a significant difference in the number of c-fos+ cells at each distance between the full experimental case (MK414-RH) and all others (all p < 0.001 at all distances for all pairwise comparisons). There was no significant difference between the glutamate-block and UOA-insertion-only cases at any distance (p > 0.23 at all distances), and both these cases differed significantly from the light-only case at 0mm distance (p < 0.05 for all comparisons). Finally, the number of c-fos+ cells decreased significantly with distance for cases MK414-RH (p < 0.001), MK422-RH (p = 0.001), and MK414-LH (p = 0.003), but not for case MK421-RH (p = 0.079).

## DISCUSSION

We have developed and validated a novel device, the UOA, which has the potential to further optogenetic research in NHPs. Current optogenetic approaches in NHPs permit light delivery either over a large superficial area^9,23^, or to deeper tissue over a small area^25-27,38^. Multi-site probes for larger volume stimulation have also been developed, and combined with single^41^ or multisite^42,43^ electrical recordings, but these approaches are typically cumbersome to assemble and don’t easily scale to precisely target multiple small tissue volumes. The UOA combines the advantages of all these approaches and retains millisecond-scale temporal resolution. It allows for both focal and larger-scale neuronal activation of single or multiple deep layers simply by varying the number of simultaneously activated μLEDs and/or the light irradiance. Moreover, although here we only used the needle-aligned μLED array for deeper layer activation, the integrated interleaved interstitial μLED array will allow for selective photostimulation of superficial layers, either independently or in conjunction with deep layers.

By design, the UOA is intended to achieve spatial resolution in cortical application in NHPs, and eventually humans, and is, thus, ideal for addressing neuroscience questions that require large-scale manipulations of deep and/or superficial cortical layers. Here we have demonstrated that the UOA, used as a stimulation-only device in conjunction with LEA recordings, can be used to study inter-laminar interactions. We were able to localize photostimulation to single or multiple cortical layers by varying light intensity. Similarly, varying insertion depth (or shank length) enables targeted selection of layers. Relative differences in onset latency of evoked responses could be used to distinguish distinct network activity patterns following different patterns of UOA stimulation. For example, at low light irradiance, direct neuronal activation was initially localized to layers nearest optrode tip termination before spreading trans-synaptically to other layers. Increasing light irradiance reduced or eliminated these latency differences. Similarly, firing rates in L4C increased less at higher versus lower intensities, suggesting response amplitude can be used to identify local activation of higher threshold inhibitory networks.

We showed that by varying the distance between the stimulation site/s on the UOA and the recording electrode, local versus long-distance intra-areal interactions can be studied. Moreover, used in conjunction with c-fos IHC, we were able to identify multi-synaptic interactions within and beyond the photostimulation area. Photostimulation via the UOA increased c-fos expression over distances much > 8mm (well beyond the stimulated cortical area), but spiking activity could not be evoked beyond ∼3 mm from the stimulated site, indicating c-fos expression revealed subthreshold activity induced by network interactions. This is consistent with previous demonstrations of c-fos expression several synapses away from an electrically stimulated site. Thus the UOA in conjunction with c-fos IHC can be used for functional mapping of neuronal circuits^39^.

We also investigated whether our results could have been affected by local increase in brain temperature caused by the μLEDs heating up when activated. This concern arises with implantable devices^44^ both in terms of temperature-induced tissue damage^45^ and changes in spiking activity^24,46^. It is generally assumed that tissue damage is negligible for temperature increases < 1°C^29,47^. One difference of the UOA compared to other implantable μLED devices is that the heat-generating μLEDs are mounted on the topside of the device and external to tissue, compensating for the fact that the low optical coupling efficiency requires higher drive currents than for optogenetic devices based upon embedded μLEDs on implantable shanks^44^. Detailed thermal simulations showed that the intervening thermally-insulating layers of dura-gel and brain tissue (combined thickness ∼1.5 mm) caused a ∼1 second delay in the temperature ramp at the stimulation site in L4, so that the bulk of the temperature rise (and subsequent fall) occurred during the inter-trial interval and not during the trial period. These simulations also showed that peak temperature rise could be held below 1°C. Additional analysis of spiking rates during the inter-trial interval showed some modulation from background activity, which could be temperature mediated, but only when the whole array was activated at the highest intensity; and even for this condition, spiking activity had returned to baseline by the end of the inter-trial interval prior to the next trial. These results strongly suggests that our results were not affected by thermal increases.

Future applications, beyond those demonstrated here, could involve functional investigations of inter-areal circuits, when UOA stimulation in one cortical area is coupled with recordings in a different area. Importantly, despite its limited shank length (∼2.5 mm maximum), the UOA can also be employed to study cortico-subcortical interactions, e.g., through modulation of axon terminals of deep nuclei within cortex, and recordings of postsynaptic cortical neurons in the same cortical area and/or layer.

In conclusion, the UOA, as currently conceived, will enable studies addressing fundamental questions in neuroscience, e.g., the role of corticocortical feedback and cortical layers in the model systems closest to humans. As many human neurological and psychiatric disorders have been linked to abnormalities in cortical circuits^4,5^, this technology can improve our understanding of the circuit-level basis of human brain disorders, and will pave the way for a new generation of precise neurological and psychiatric therapeutic interventions via cell type-specific optical neural control prosthetics.

## ONLINE METHODS

### Device Fabrication, Characterization, and Benchmarking

Fabrication and testing of the first generation UOA devices was previously reported^28,48^. The second-generation devices used in this study included an optical interposer layer that limits emission from the μLED array to the shank sites for illumination of deep cortical tissue.

### Fabrication

A 2 mm-thick, 100mm diameter Schott Borofloat 33 glass wafer used to construct the optrode needles was anodically bonded to a freshly cleaned 0.1mm thick, 100 mm diameter intrinsic Si wafer serving as an optical interposer. The Si and Borofloat wafers were coarsely aligned, and bonding performed using an EVG 520 anodic bonder. The optical vias were patterned in the Si interposer by deep reactive ion etching (DRIE) using a Bosch process. A 10-μm-thick AZ9260 soft mask was photolithographically patterned to define the array of 80×80 μm^2^ optical vias for shank and interstitial illumination for the DRIE process. The bonded wafer was then sub-diced into *modules* of 9 to 16 UOAs using a DISCO 3220 dicing saw.

UOA modules were mounted to a carrier wafer using WaferGrip™ (Dynatex International, Santa Rosa, CA). The glass shanks were cut with the DISCO 3220 using the previously reported process^28,48^. Briefly, beveled blades were first used to generate pyramidal tips on the surface, followed by standard profile blades to form the shanks. The shanks on a module were then etched to a nominal 110μm thickness using a mixture of hydrofluoric (49%) and hydrochloric (37%) acid in a 9:1 ratio. The die was then demounted and cleaned, and the shanks were smoothened to decrease light scattering using a 725 °C heat treatment for 2 hours in a vacuum furnace. UOA modules were then singulated into individual 4×4 mm^2^ UOAs using the DISCO 3220.

Arrays of μLEDs on thinned (150μm) sapphire substrates, from the Institute of Photonics at University of Strathclyde, were integrated with the UOA using closed-loop optical alignment to the optical vias on individual UOAs at Fraunhofer IZM (Berlin, Germany)^28^, and bonded using index-matched epoxy. At the University of Utah, passive matrix μLED pads were wire bonded to an ICS-96 connector (Blackrock Microsystems, Salt Lake City, UT) using insulated gold alloy wire. The wire bundle and backside of the UOA were then potted in NuSil MED-4211 silicone, respectively, followed by overcoating with a 6μm-layer of Parylene C.

### Bench Testing

To characterize the electrical and optical performance of the finalized devices, the latter were attached to a custom switchboard for matrix addressing the individual optrode shanks. The switchboard consisted of a matrix arrangement of parallel-connected mechanical switches and electrical relays, 10 sets for the anodes and 10 sets for the cathodes.

This enabled both manual and automated activation of individual optrode shanks or optrode patterns. For the automated activation and testing, the relays were connected to Arduino boards that received commands from the lab computer. To prevent voltage spikes originating from the switching of the channels from damaging the μLEDs, the anode paths also contained a small filter circuit consisting of capacitors and Zehner diodes (break-down voltage: 8.2V). For the automated testing, the UOAs were inserted into the opening of an integrating sphere that was, in turn, connected to a photodetector and power meter (Newport 2832-C Dual-Channel Power Meter). The calibration factor of the integrating sphere was determined using a fiber coupled LED prior to the experiment. Then the UOAs were connected to the switchboard, and the latter was connected to a source measure unit (Keithley 236 Source Measure Unit) for the measurement. The automated characterization was conducted as follows: the switchboard’s Arduino boards received the command to switch to an individual optrode shank using the relays. Then the source measure unit applied a voltage pulse measurement pattern (pulse length 100ms, pause between pulses 1900ms to prevent heat buildup) sweeping the voltage from 0 to 7.2V (or until the compliance current of 100mA was reached) with each pulse increasing by 100mV. For each pulse, the resulting current and the output optical power were recorded; the optical power was then corrected using the integrating sphere calibration factor. This was repeated for each individual optrode shank of the device for a full characterization.

To ensure the stability of the device for an acute *in vivo* experiment, additional voltage transient measurements were made before and after a 48-hour soak test in phosphate-buffered saline (PBS) at 37 °C. Further, an electrode was immersed in solution to verify encapsulation integrity, as evidenced by lack of shorting to solution.

For the *in vivo* experiments, the switchboard was upgraded two-fold: first, transistors were added to the cathode channels to allow for turning the device on and off based on an external TTL trigger. However, we found that turning on the optrodes using the trigger signal directly induced too strong a capacitively-coupled voltage signal in the recording. Therefore, as a second upgrade, an additional Arduino board with digital-analog-converter was added that received the external trigger and introduced rise and fall times to the square wave. This reduced the capacitively-coupled interference to a level below measurable when both the LEA and the UOA were in close proximity in 1xPBS solution prior to the *in vivo* experiment. During the experiment, the voltage for the UOA was supplied by a lab power supply via the switchboard, and the switches were operated manually to define the required patterns.

### Modeling

To understand light spread in tissue, the optical output of the device was modeled using ray-tracing software (Optics Studio 12, in non-sequential mode). This model has been described previously^28^. Brain tissue was modeled using a Henyey-Greenstein scattering model, with a scattering coefficient of 10 mm^-1^, absorption coefficient of 0.07 mm^-1^, and anisotropy of 0.88^47^. Each needle was modelled individually using its measured optical output at the given voltage level. To generate the cross-section images from a simultaneously illuminated column (**Fig. 1G**), the light output from the 10 needles in that column were summed.

### Animals

A total of 3 adult female Cynomolgus monkeys (*Macaca fascicularis*) were used in this study. The left hemisphere of one animal (case MK421-LH) was used for the *in vivo* electrophysiological testing of the active UOA (integrated with the μLED array). The right hemisphere from the same animal (MK42-RH), and 3 hemispheres from 2 additional animals (MK414RH and LH, and MK422-RH) were used for c-fos testing of the passive UOA (i.e., without an integrated μLED array). All procedures conformed to the National Institutes of Health Guide for the Care and Use of Laboratory Animals and were approved by the University of Utah Institutional Animal Care and Use Committee.

### Survival Surgical Procedures and Viral Injections

Animals were pre-anesthetized with ketamine (10□mg/kg, i.m.), intubated, placed in a stereotaxic apparatus, and artificially ventilated. Anesthesia was maintained with isoflurane (1– 2.5% in 100% oxygen). Heart rate, end tidal CO2, oxygen saturation, electrocardiogram, and body temperature were monitored continuously. I.V. fluids were delivered at a rate of 3-5/cc/kg/hr. The scalp was incised and a craniotomy and durotomy were performed over area V1 (n=2 animals, MK421-LH and MK414-RH), or rostral to the precentral gyrus, roughly above the supplementary motor area (SMA; n=1, MK422-RH). We injected a 1:1 viral mixture of AAV9.CamKII.4.Cre.SV40 and AAV9.CAG.Flex.ChR2.tdTomato (Addgene Catalog #s: 105558, and 18917, respectively). We have previously found that this method nearly eliminates retrograde expression of transgenes^9^. The viral mixture was slowly (∼15nl/min) pressure-injected (250-350nl repeated at 2 or 3 cortical depths between 0.5 and 1.5 mm from the cortical surface) using a picospritzer (World Precision Instruments, FL, USA) and glass micropipettes (35-45μm tip diameter). After each injection, the pipette was left in place for 5-10 min before retracting, to avoid backflow of solution. A total of 5-6 such injections, each 500-750nl in total volume, and spaced 1.5-2mm apart, were made in two animals (MK421-LH, MK414-RH) while the third animal (MK422-RH) received 2 × 1,050nl injections. These injections resulted in a region of high viral expression roughly 4-6 mm in diameter (as an example see **Extended Data Fig. 3A Right**). Following viral injections, a sterile silicone artificial dura was placed on the cortex, the native dura was sutured and glued onto the artificial dura, covered with Gelfoam to fill the craniotomy, and the latter was sealed with sterile parafilm and dental acrylic. Anesthesia was discontinued and the animal returned to its home cage. After a survival period of 5-10 weeks, to allow for robust ChR2 expression, the animals were prepared for a terminal UOA photostimulation procedure.

### Terminal Surgical Procedures and UOA Insertion

Monkeys were pre-anesthetized and prepared for experiments as described above. Anesthesia and paralysis were maintained by continuous infusion of sufentanil citrate (5–10□ μg/kg/h) and vecuronium bromide (0.3□mg/kg/h), respectively. Vital signs were continuously monitored for the duration of the experiment, as described above. Following suture removal and scalp incision, the craniotomy and durotomy were enlarged to allow space for device implantation, and ChR2 expression was verified *in* vivo using a custom fluorescent surgical microscope (Carl Zeiss, GmbH; **Fig. 2B**). UOAs were positioned over cortical regions of high tdT/ChR2 expression (e.g. **Figs. 2B, 6B, F**), and then inserted using a high speed pneumatic hammer typically used for insertion of Utah Electrode Arrays^34^ (Blackrock MicroSystems, UT). Parameters used for insertion were 20 psi for 30ms, using a 1 mm-long inserter, in order to achieve partial insertion of the UOA, so as to minimize insertion trauma on the cortex. In two animals used for c-fos experiments after partial insertion with the pneumatic inserter, the UOA was gently pushed down to achieve deeper insertion.

### Photostimulation

We implanted two types of UOA devices: (i) a 10×10 UOA with fully integrated μLED arrays (also referred to as “active” device; n=1 device in 1 animal, MK421-LH; see **Fig. 2A-C**), and (ii): 10×10 UOAs with an optical interposer integrated into the sapphire backplane, but with no μLED array for light delivery (referred to as “passive” devices; n=3 devices in 3 hemispheres from 2 animals, MK414-RH, MK414-LH, MK422-RH). The active device was used for electrophysiological testing experiments, while the passive devices were used for the c-fos experiments.

### Active Device (Electrophysiology)

Photostimulation with the active UOA occurred via the integrated ***Active Device*** LED array. Photostimulation parameters were 5Hz, 100 msec-pulse duration for 1 sec, followed by 1.5-21sec inter-trial interval (longer intervals were used at the higher photostimulation intensities). We varied the spatial pattern (single μ LED along column 1, entire single columns, and all μ LEDs across the entire UOA) and intensity (from 2.8 to 7.8V input intensity) of photostimulation as described in the Results section.

### Passive Devices (c-Fos)

Selective photostimulation via passive devices was obtained by illuminating a subset of UOA needles with an appropriately positioned fiber-coupled 473nm laser (400 μ m multimode optic fiber, ThorLabs Newton, NJ; laser: Laserwave, Beijing, China) held in place with a stereotaxic tower. We used a collimating lens (ThorLabs, Newton, NJ) to restrict spot size to ∼ 1.5mm in diameter. To shield stray light, we covered any exposed tissue around the illuminated area, as well as the non-illuminated portions of the UOA, with an opaque (black) artificial dura. For each UOA we stimulated 2 or 3 separate sites. At each site we used phasic photostimulation (50Hz for 2.5 min, 2.5 min pause, and 20Hz for an additional 2.5 min; pulse duration was 10ms) at 3.8mW power output (corresponding to an estimated irradiance of 15-19mW/mm^2^).

### Electrophysiological Recordings

Extracellular recordings were made in V1 with 24-channel linear electrode arrays (LEAs; V-Probe, Plexon, Dallas, TX; 100μm contact spacing, 300μm from tip to first contact, 20μm contact diameter). The LEAs were inserted into the cortex next to the UOA to a depth of 2.4-2.6mm, slightly angled laterally (towards the UOA) and posteriorly. We made a total of 3 penetrations (P1-P3; **Extended Data Fig. 3A**), of which only P2 and P3 provided useful data. After UOA and LEA were inserted into the cortex, we applied a layer of Dura-Gel (CambridgeNeuroTech, Cambridge, UK) over the cortex and UOA, to prevent the cortex from drying and stabilize the recordings. A 128-channel recording system (Cerebus, Blackrock Microsystems, Salt Lake City, UT) was used for standard signal amplification and filtering. Multi-unit spiking activity was defined as any signal deflection that exceeded a voltage threshold (set at 4 x the SD of the signal on each channel). Threshold crossings were timestamped with sub-millisecond accuracy. We did not record responses to visual stimuli but only to UOA photostimulation performed as described above; thus, the monkey’s eyes were closed during the duration of the experiment.

### Analysis of Electrophysiological Data

We analyzed MUA responses from a total of 45 contacts deemed to lie within the parafoveal representation of V1 in two penetrations (out of 3 total, see above) for which neural activity was modulated by photostimulation via the active UOA. For the results presented in **Figures 3-5**, quantitative analysis was limited to contacts on which MUA was stimulus modulated (one-way ANOVA comparing spike rates during full one-second photostimulation trials with spike rates during control periods of equivalent duration, p < 0.01).

To quantify the change in MUA firing rates, relative to background, during photostimulation we calculated firing rates for all pulse epochs within all trials and then compared them to the average background rate. To estimate the preference at each recording site for stimulation across the full range of tested UOA locations (**Fig. 3**), we regressed average evoked-responses on UOA stimulation site and intensity. Preliminary analyses had revealed a non-monotonic relationship between stimulation intensity and response on many contacts (cf. **Fig. 2F**), thus we included a quadratic term in the regression model.

### CSD analysis

For the CSD analysis shown in **Fig. 2D-E**, current source density (CSD) was calculated from the band-pass filtered (1-100Hz) and pulse-aligned and averaged LFP, using the kernel CSD toolbox (kCSD Matlab)^49^. CSD was calculated as the second spatial derivative of the LFP signal, reflecting the net local transmembrane currents generating the LFP. The depth profile of the CSD was estimated by interpolating every 10μm. To facilitate comparisons across conditions, CSDs from different conditions were normalized to the standard deviation (SD) of the baseline (50ms prior to pulse onset) after subtraction of the baseline mean.

### Onset Latency

To quantify the onset latency of MUA responses, we either: (i) calculated the average peri-stimulus time histogram (PSTH) from all pulse-aligned responses (e.g., **Fig. 4**) or (ii) estimated a PSTH separately for the response to each pulse (**e.g**., **Extended Data Fig. 7**). Peristimulus time histograms (PSTHs) were estimated via an adaptive algorithm in which the MUA raster was first convolved with a Gaussian kernel of fixed width (3ms bandwidth), kernel width was then adapted so that the number of spikes falling under the kernel was the same on average across the response (http://chronux.org^50^). We then subtracted the mean baseline response from the stimulus-evoked response. For each response measure, i.e., either the average or pulse-by-pulse PSTHs, we took the time at which the response reached 25% of the peak as the onset latency (results were qualitatively similar using 15% and 35% criteria; data not shown). We report the former measure as the mean onset latency in **Figures 4-5**. We used the latter measure to test for differences in onset latency across contacts within and across UOA stimulation parameters (**Figs. 4-5 and Extended Data Fig. 7**).

### Statistical Analysis

Stimulus-evoked firing rates were calculated from pulse-aligned or trial-aligned responses and baseline corrected (mean baseline activity subtracted). We determined responsiveness to stimulation via a one-way ANOVA comparing firing rates during the full 1-second trial period with inter-leaved control periods of equivalent duration; MUA at an LEA recording site was deemed responsive if there was a significant difference between stimulation and control trials at the p=0.01 level. To estimate the selectivity of MUA for stimulation at different UOA sites we performed a multiple linear regression, with UOA column, row, and intensity as independent variables and pulse-aligned, baseline corrected, firing rates as the dependent measure. To test for differences in the goodness-of-fit of models with- and without a quadratic term, we used a two-sample Kolmogorov-Smirnov test. We assessed the effects of varying UOA stimulation site and intensity on response amplitude or onset latency using ANOVA models followed by the Tukey-Kramer test for post-hoc comparisons.

### c-Fos Experiments

We used 4 hemispheres from 3 animals for these experiments (MK414-RH and LH, MK422-RH, and MK421-RH). Two of these animals (MK422 and MK414) were prepared for a terminal experiment (as described above) 5 or 10 weeks, respectively, after the viral injections, and a passive UOA was inserted in regions of high tdT/ChR2 expression in the injected hemisphere. In one of these animals (MK422-RH), UOA insertion was preceded by glutamate block (see below). After UOA insertion, photostimulation was performed via an optical fiber-coupled laser through the UOA, as described above. Two additional hemispheres in 2 animals (MK414-LH and MK421-RH) were used as controls. Specifically, case MK414-LH received insertion of a passive UOA in non-opsin expressing SMA cortex, and was euthanized 4 hours following UOA insertion without receiving any photostimulation. As a separate control, in case MK421-RH we performed surface photostimulation of SMA cortex not expressing opsins, using a fiber-coupled laser and a collimating lens and the same photostimulation protocol described above for other c-fos experiments; no UOA was inserted in this case. In all animals, UOA insertion and/or photostimulation were performed after a 10-14-hour period of eye closure and at least 5 hours after completion of surgery, and the animals were euthanized 75 minutes after completion of the photostimulation protocol.

### Pharmacological Blockade of Local Glutamate Signaling

To compare changes in c-fos expression due to direct local optogenetic activation with indirect local and long-range changes due to synaptic increases in excitatory glutamatergic neurotransmission downstream of the directly-activated neurons, in one case (MK422-RH) we applied the selective glutamate AMPA receptor antagonist 2,3-dihydroxy-6-nitro-7-sulfamoyl-benzoquinoxaline-2,3-dione (NBQX, 5mM) (Tocris BioSciences). NBQX was applied topically prior to UOA insertion, by soaking a piece of Gelfoam placed over ChR2-expressing SMA cortex with 1ml of the drug solution. The drug was allowed to passively diffuse through the cortical layers for 90 minutes, during which 100-200μl of the solution were applied every 15 minutes to ensure saturation of the Gelfoam, after which the Gelfoam was removed and the passive UOA inserted over the region of glutamate block. Photostimulation was performed as described above for the passive device.

### Histology

On completion of the experiments, the animals were euthanized by an overdose of Beuthanasia (0.22 ml/kg, i.v.) and perfused transcardially with saline for 2–3 min, followed by 4% paraformaldehyde (PFA) in 0.1M phosphate buffer for 20 min to fix the brain. The brains were post-fixed overnight in the same fixative, sunk in cryoprotectant 30% sucrose solution, and sectioned at 40μm on a freezing microtome. The hemisphere used for electrophysiological testing of the active UOA (MK421-LH) was sectioned tangentially. One in 3 sections were wet-mounted and imaged for fluorescent tdT-label at 10x magnification. The same sections were then reacted for cytochrome oxidase (CO) to reveal cortical layers and the location of UOA and LEA insertions visible as discolorations in CO staining (**Extended Data Fig. 3A Left**).

All other hemispheres used for c-fos experiments were sectioned sagittally. One full series of sections (1:3) were immunoreacted for c-fos by overnight incubation in primary antibody (1:500 rabbit anti-c-fos, Ab 19089, Abcam, MA, USA) at room temperature, followed by 2 hrs incubation in near-infrared secondary antibody (1:200 donkey anti-rabbit IgG-AF647, Jackson ImmunoResearch, PA, USA) at room temperature. Sections were then wet-mounted, counterstained with blue fluorescent Nissl (1:100 N21479, Thermo Fisher Scientific, MA, USA) by dripping the solution onto the slide-mounted sections every 5 min for 20 min, rinsed, and coverslipped and sealed with CoverGrip™ Coverslip Sealant (Biodium, CA, USA).

### Tissue Imaging

Imaging of tissue sections was performed on a Zeiss Axio Imager.Z2 fluorescent microscope (Zeiss, Germany) with a Zeiss X-cite 120 LED Boost light source, using a 10x objective and an Axiocam 506 mono camera (Zeiss, Germany). Image files were created and analyzed using Zen 2.6 Blue Software (Zeiss, Germany). The light intensity was set to 100%, and the exposure time for each channel was kept the same between images. The tangentially-sectioned hemisphere (MK421-LH) was imaged as described above. In all other cases, each sagittal section was imaged in 3 channels simultaneously, one channel for tdT/ChR2 (red-but note the color was artificially changed to green in **Fig. 6B, F**), one channel for Alexa-647-c-Fos (far-red), and the third channel for 435-455 Nissl (blue).

### Analysis of c-Fos Expression

To quantify c-fos expression, c-fos+ cells were plotted and counted in sampled areas, using Neurolucida software 2006 (Microbrightfield Bioscience, VT, USA). For each case, we selected for counts 5 sections spaced 1 mm apart encompassing the area of UOA insertion and/or photostimulation (for the light-only case). In each section, we plotted and counted cells within three 200μm-wide windows spanning all cortical layers, one positioned at or near the center of the UOA insertion region (or of phtostimulation-only), and the other two located at distances of 4mm and 8mm, respectively, from the center of the UO insertion (**Fig. 6**). Thus, a total of 15 regions of interest (ROIs) were counted for each case. The laminar distribution of c-fos+ cells was analyzed by tracing the layers on the Nissl stain and counting the number of c-fos+ cells within each layer in Neurolucida. Statistical differences in c-fos+ cell counts among experimental and control cases, and across distances were estimated using a one-way ANOVA with Bonferroni corrected post hoc comparisons.

## DATA AVAILABILITY STATEMENT

The data presented here will be provided upon reasonable request to the corresponding authors.

## CODE AVAILABILITY STATEMENT

All custom software used for analysis and modeling will be provided upon reasonable request to the corresponding authors.

## ACKNOWLEDGMENTS

We thank Kesi Sainsbury for histological assistance, Seminare Ta’afua for help with experiments, and Julian Haberland and Christine Kallmayer at the Fraunhofer-Institut für Zuverlässigkeit und Mikrointegration (IZM) for □LED-to-interposer bonding. This work was supported primarily by a BRAIN grant (U01 NS099702) from the National Institute of Health (NIH) to S.B., A.A. L.R and S.M. Additional support was provided by grants from the NIH (R01 EY026812, R01 EY019743, R01 EY031959), the National Science Foundation (IOS 1755431), and the Mary Boesche endowed Professorship, to A.A, and an unrestricted grant from Research to Prevent Blindness, Inc. and a core grant from the NIH (EY014800) to the Department of Ophthalmology, University of Utah.

## AUTHOR CONTRIBUTIONS

Conceptualization: A.C., A.I., C.F.R., N.M., L. R., K.M., S.B. A.A. Device Fabrication and Bench Testing: C.F.R., Y.C., N.M., L.R.,K.M., M.D.D., S.B. Modeling: N.M., K.M. *In Vivo* Electrophysiology Testing: A.C., D.C., C.F.R., F.F., A.A. *In Vivo* cFos Testing: A.I., J.B., F.F., J.D.R., A.A., A.C. Analysis of Electrophysiology Data: A.C. Analysis of cFos Data: A.I., J.B., F.F., A.A. Writing-Original Draft: A.C., L.R., S.B., A.A. Writing–Review/Editing: all authors. Supervision & Funding Acquisition: A.A., S.B., K.M., L.R.

