## Supplemental Figure 1 for "An Optrode Array for Spatiotemporally Precise Large-Scale Optogenetic Stimulation of Deep Cortical Layers in Non-human Primates"

#### A UOA output power (mW) at different input voltages

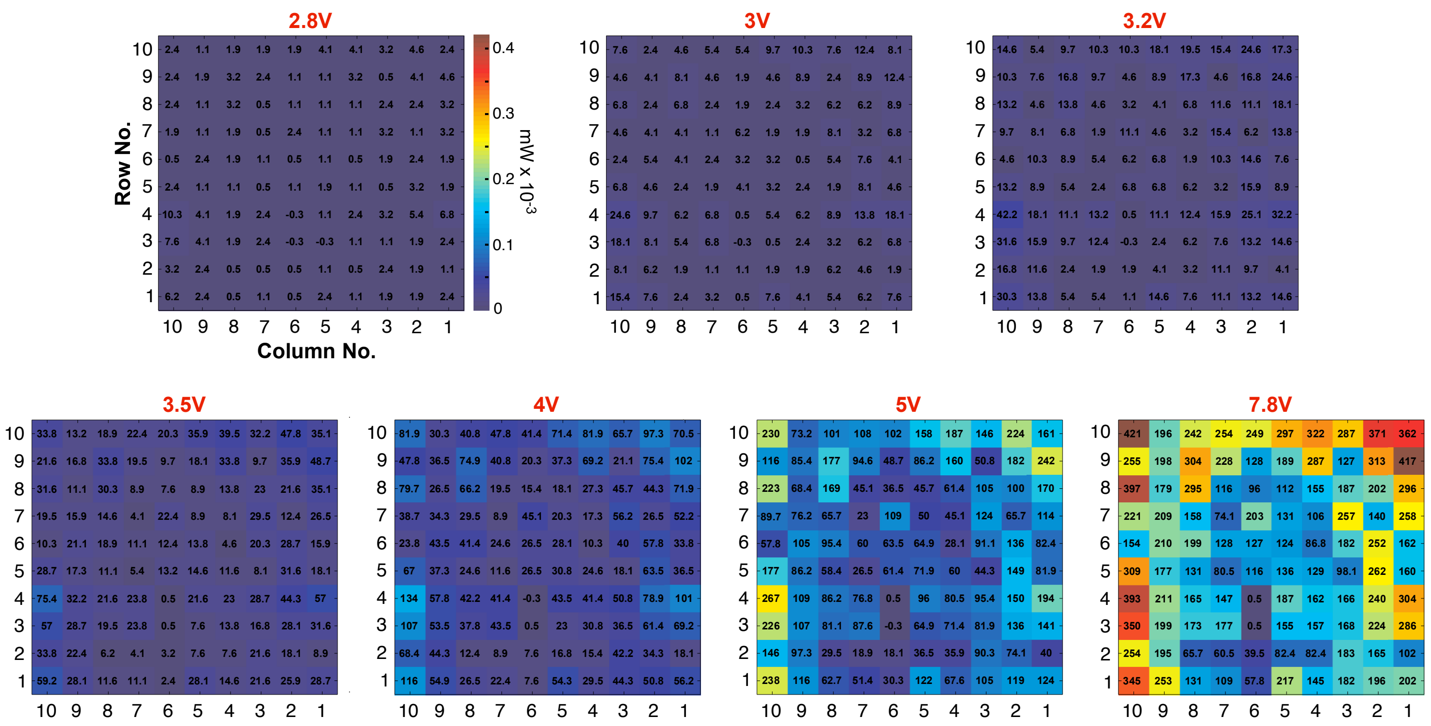


#### B UOA output irradiance (mW/mm2) at different input voltages

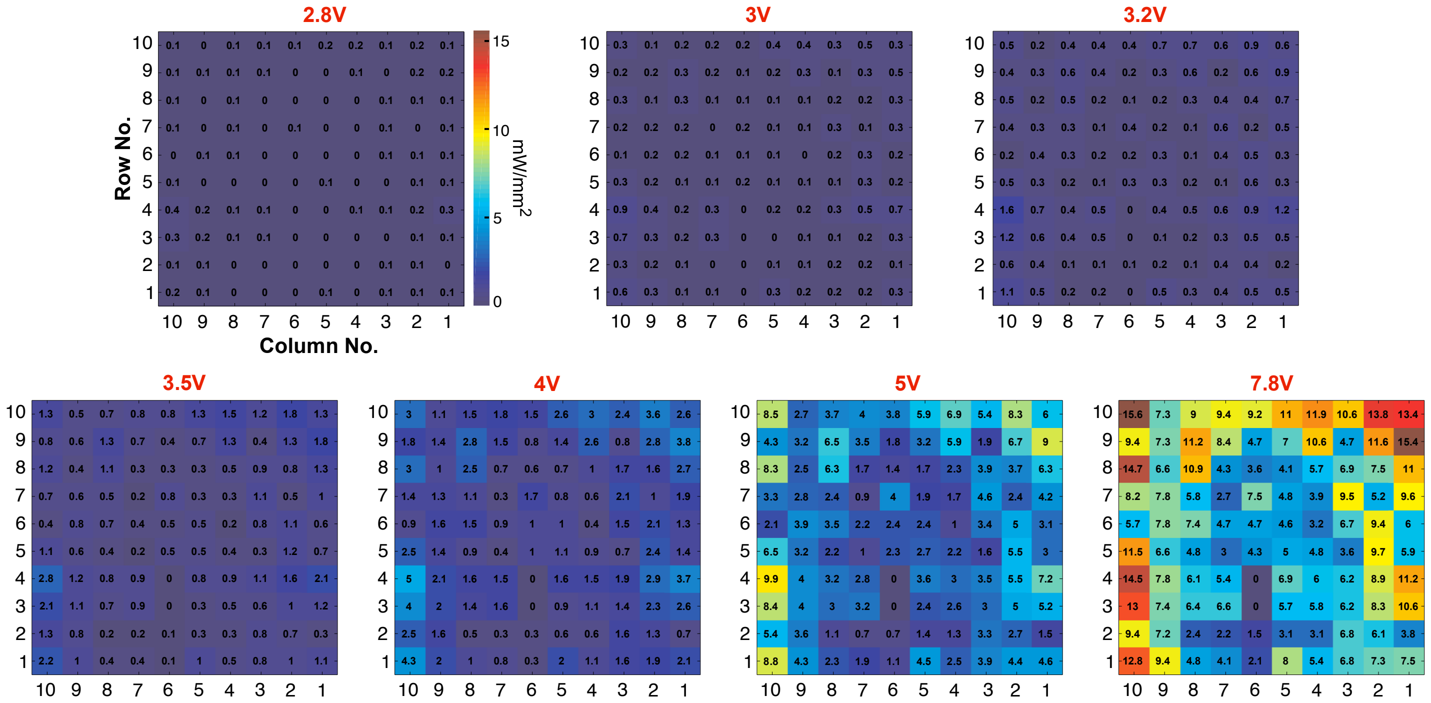


### Extended Data Figure 1

**Output optical power and irradiance of the UOA**

Each heat map represents the output optical power **(A)** or irradiance **(B)** measured at each needle tip across the entire UOA for different input voltages (indicated at the top of each map). We defined the irradiance as the emitted optical power divided by the area of the emission surface. Needles C6-R3 and C6-R4 did not emit light at their tips, thus, they were not included in the descriptive statistics in Extended Data Table 1.
