## Supplemental Figure 2 for "An Optrode Array for Spatiotemporally Precise Large-Scale Optogenetic Stimulation of Deep Cortical Layers in Non-human Primates"

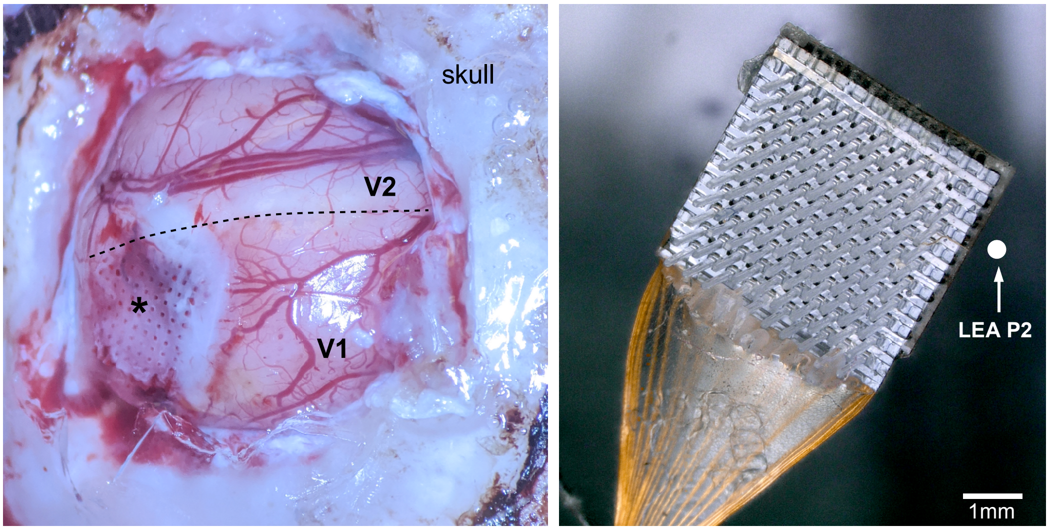


**Extended Data Figure 2**

**UOA after explantation**
**(A)** Image of the brain after explantation of the UOA at the end of the *in vivo* testing experiment. The *asterisk* marks the center of the UOA implantation site. The white area at the site of the explantation is the *Duragel* that was placed on the brain after insertion of the UOA to protect the cortical surface and prevent deydration. The *dashed line* marks the border between V2 and V1. **(B)** The UOA after explantation. The *white dot* indicates the approximate location of the LEA used for penetration 2 (P2) relative to the UOA. The shank at the top right corner (column 1 row 10) was broken prior to the UOA insertion. The remaining shanks are intact.
