## Supplemental Figure 3 for "An Optrode Array for Spatiotemporally Precise Large-Scale Optogenetic Stimulation of Deep Cortical Layers in Non-human Primates"

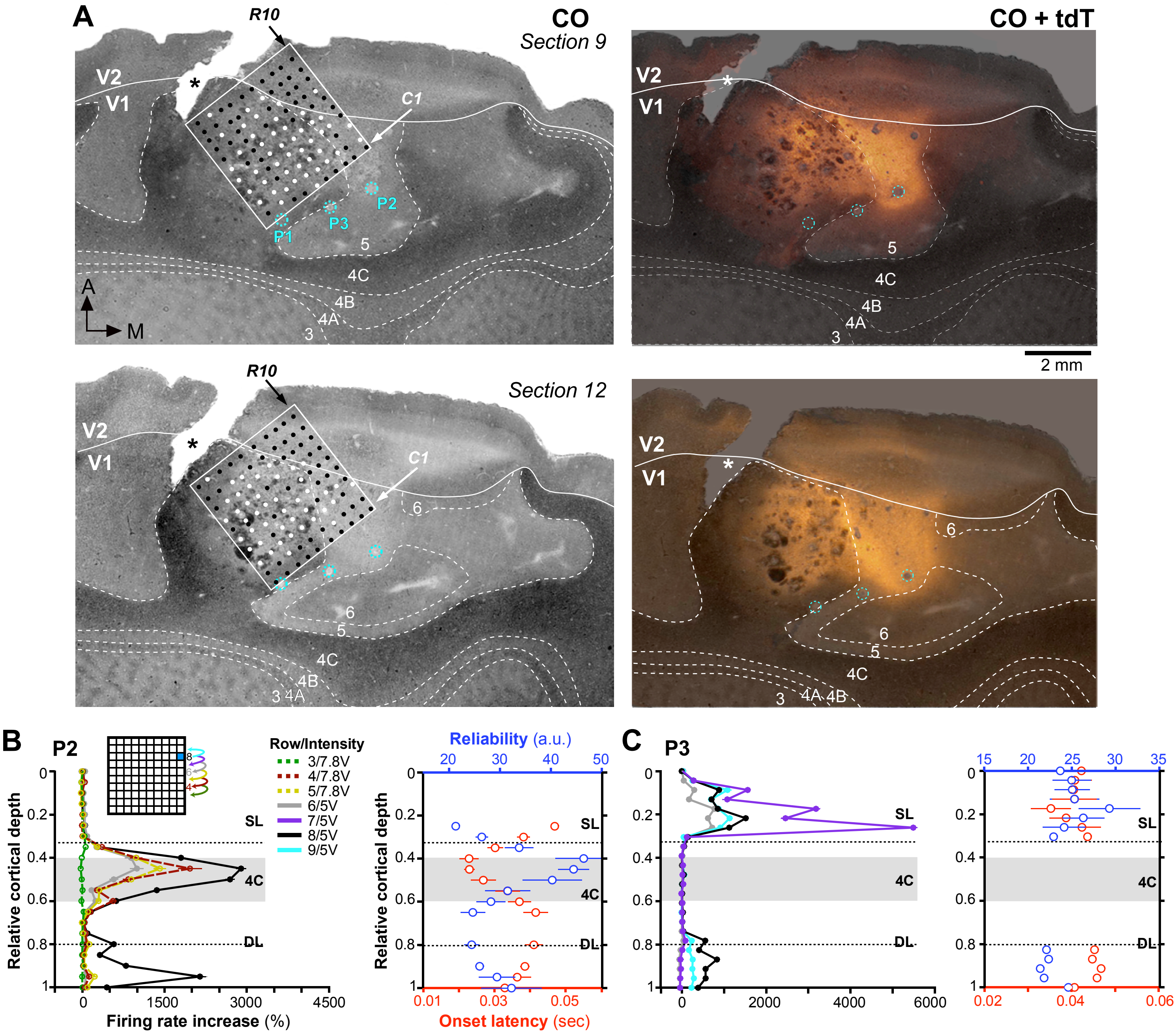


### Extended Data Figure 3

#### Laminar and tangential location of UOA and LEA penetrations

**(A)** **Left:** The locations of the UOA (*white box*) and of 3 LEA penetrations *(P1-P3, cyan dashed circles*) are shown on two cytochrome-oxidase (CO)-stained tangential sections through V1 and V2 (top section is more superficial). *Solid white contour*: V1/V2 border; *dashed white contours* delineate V1 layers (indicated). The location of column 1 (*C1*) and row 10 (*R10*) are indicated by *arrows*. *White dots* mark the locations of UOA needles visible in these sections as cortical damage. *Black dots* mark the location of UOA needles visible in more superficial sections, but not in these sections. Note that the majority of white dots are located in layer 4C in both sections. There are no white dots in column 1 in section #12 and only 2 in section #9, as the needle tips in this column terminated in the sections just above, thus in the superficial part of L4C. The postero-lateral half of the UOA terminated in slightly more superficial layers compared to its anterio-medial half. *A:*anterior; *M:* medial. **Right:** Same CO-stained section as shown on the left, with superimposed image of the same section viewed under tdT fluorescence. The fluorescent image was rendered transparent in Adobe Photoshop. P2 and P3 were located inside or near, respectively, the region of tdT/ChR2 expression, whereas P1 was more distant from it; accordingly, only the neurons recorded in P2 and P3, but not in P1, could be modulated by the UOA. P2 was located about 1-1.1 mm medial to the nearest UOA needle (C1-R8, C1-R9, in these sections), while P3 was located about 800µm from the UOA (C1-R5 and C1-R4 are the nearest needles to P3 in these sections, but, as this penetration was not vertical, the more superficial LEA contacts were closer to needles C1-R6 and C1-R7 and more distant from the UOA, as also indicated by the physiological recordings in (C), and in **Fig. 3F**). The *asterisk* in all panels marks a crack in the tissue caused by histological processing, not by the UOA insertion. **(B)** **Left:** Relative cortical depth of each contact on the LEA in P2 is plotted versus the increase in firing rate caused by stimulation of single µLEDs along column 1 (*inset*). Different color traces are data for different µLEDs (rows 3-9) at 5 or 7.8V stimulation intensity (the most distant µLEDs only evoked responses at the higher intensity). µLED C1-R8 evoked the max response, indicating this needle tip was the closest to P2. **Right:** Relative cortical depth on the LEA-P2 is plotted versus the mean onset latency (*red*) or the mean onset latency reliability (*blue*; inverse of the SD of the distribution of pulse by pulse onset latencies) of responses at each contact evoked by stimulation of the whole µLED array; means are averages across all photostimulation intensities ≤5V. The shortest and most reliable response latencies are for contacts in L4C, indicating the UOA tips nearest P2 ended in this layer. **(C)** Same as in (B) but for P3. The data indicate that the µLED closest to P3 was C1-R7 whose tip terminated in the superficial layers.
