## Supplemental Figure 4 for "An Optrode Array for Spatiotemporally Precise Large-Scale Optogenetic Stimulation of Deep Cortical Layers in Non-human Primates"

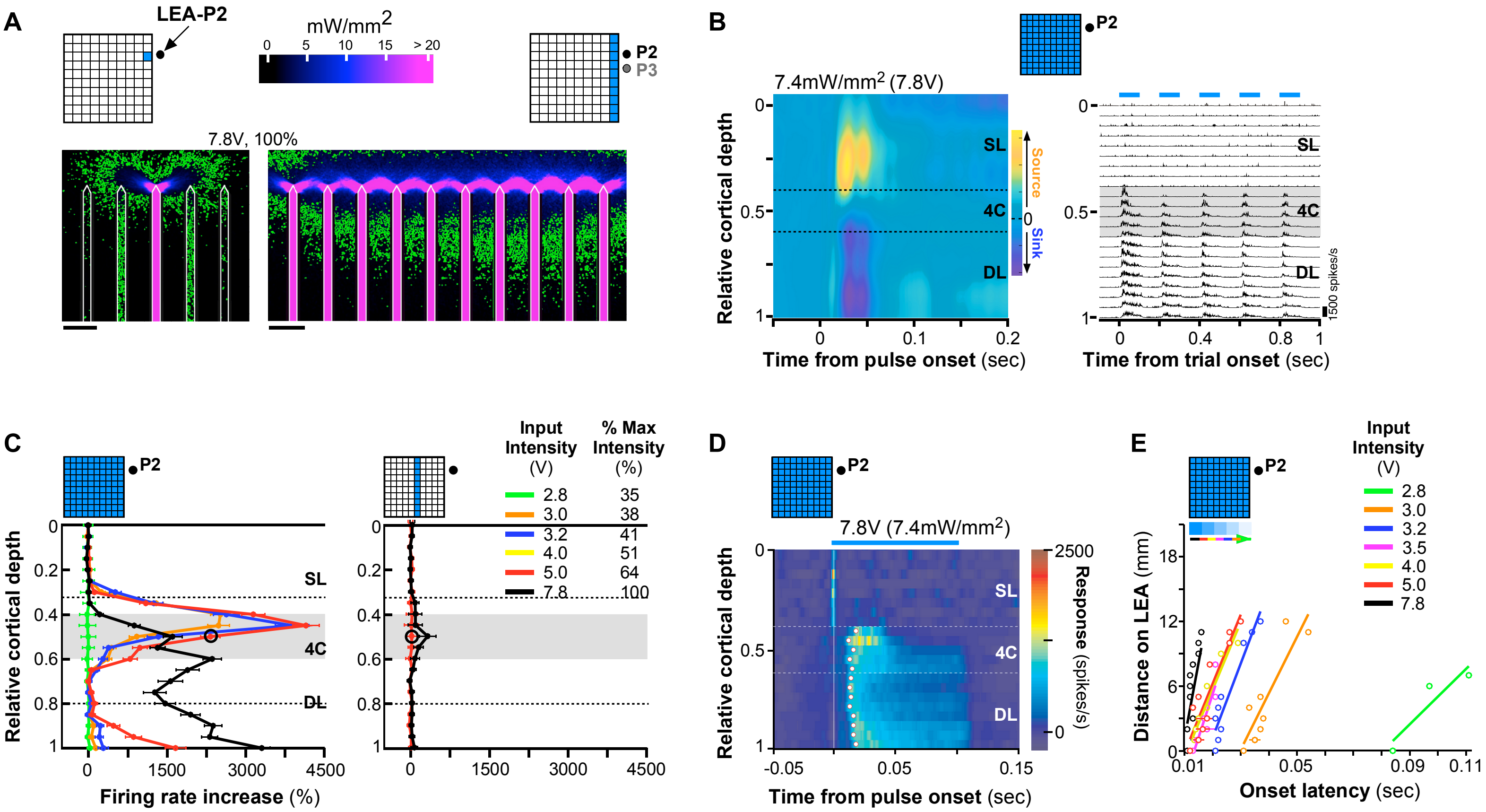


### Extended Data Figure 4

**Bench testing and *in vivo* results for UOA stimulation intensity of 7.8V**

**(A)** **Left:** Ray trace model of light spread in cortical tissue when a single µLED (C1R8, i.e. the closest to P2) is activated at 7.8V. **Right:** Model of light spread in tissue when all of column 1 (nearest to both P2 and P3) is activated at 7.8V. Green contour encloses tissue volume within which the light irradiance is above 1mW/mm$^2$. Scale bars: 400µm. **(B)** CSD analysis (**Left**) and MUA (**Right**) through the depth of V1 for P2 in response to phasic UOA photostimulation (100ms pulse, 5Hz, 7.4mW/mm$^2$) of the entire UOA. Conventions are as in **Fig. 2D**. Current sinks and strong phasic MUA in response to UOA stimulation extended from L4C to the white matter boundary. **(C)** Relative cortical depth of contacts on P2 plotted versus firing rate increase in response to stimulation of the whole UOA (**Left**) or Column 5 (**Right**) at different intensities (different color traces). Same plots as shown in **Fig. 2F and I**, respectively, with added data from 7.8V stimulation. **Left:** Relative to stimulation at lower intensities, 7.8V stimulation of the whole UOA resulted in a decreased activation peak in L4C and increased responses in L5-6. At this higher intensity, light spread through the cortical depth may have directly contributed to firing rate increase in L5 and 6, while the reduced peak in L4C may have resulted from activation of higher threshold inhibitory networks. **Right:** Stimulation of Column 5 evoked a response in L4C only at the highest intensity used (7.8V). **(D)** Heatmap of MUA through the depth of V1 during the peri-pulse period, when the whole UOA was stimulated at 7.8V. Other conventions are as in **Fig. 4**. There was no difference in onset latency of evoked MUA across layers suggesting light spreading deep into tissue at this intensity directly activated both mid- and deep layers. **(E)** Distance on the LEA of each contact from the contact with the fastest onset latency plotted against onset latency, for the whole UOA condition. Same as in **Fig. 5A** with added data from 7.8V stimulation.
