## Supplemental Figure 5 for "An Optrode Array for Spatiotemporally Precise Large-Scale Optogenetic Stimulation of Deep Cortical Layers in Non-human Primates"

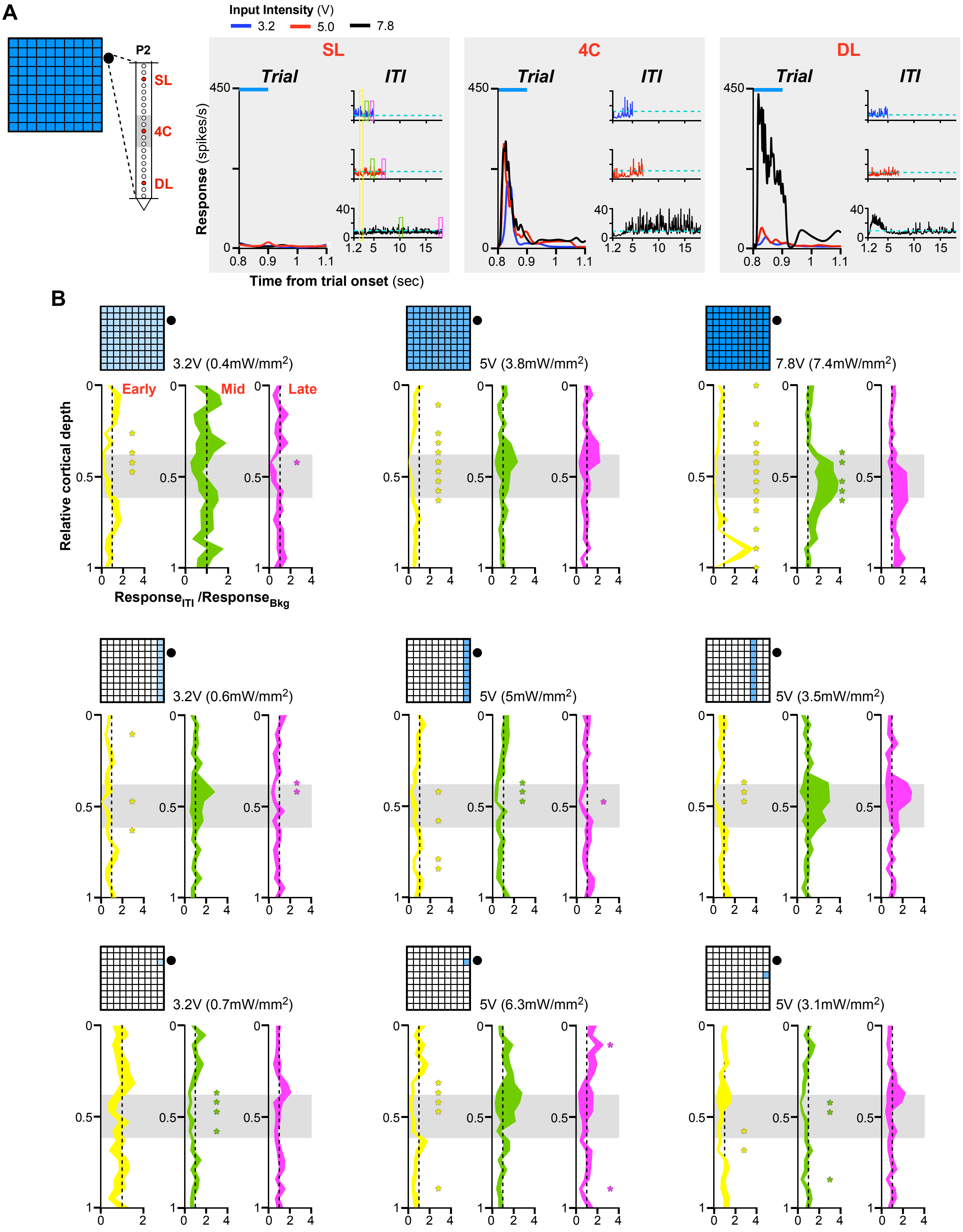


**Extended Data Figure 5**

**Neuronal activity during the inter-trial period.**

**(A)** **Left**: Schematics of whole array UOA stimulation and of LEA in P2. Data from the 3 highlighted LEA contacts in superficial layers (SL), L4C, and deep layers (DL), respectively, are shown on the right. **Right**: In each panel, trial-aligned PSTHs for the last portion of the trial period (**Left**) and the inter-trial interval (ITI; **Right**) are shown for the highlighted channels at three intensities of whole array stimulation. Each trial consisted of 1sec of photostimulation (100msec pulse duration, 5Hz) followed by 1.5-21sec ITI. *Blue bar*: final 100ms stimulus pulse period. The *yellow* (early), *green* (middle), and *magenta* (late) *boxes* in the ITI plots for the example SL channel indicate the portions of the ITI that were compared with the average ongoing activity on each channel prior to the beginning of the experiment ("background"). *Dashed cyan lines*: average MUA spike rate during a brief background period prior to each experiment. For the example contacts, there was an increase in MUA activity, relative to the average background rate, during the ITI for the 7.8V intensity in L4C and the early ITI period in the DL. **(B)** Comparison of early, middle, and late ITI period activity with ongoing background activity. For each condition shown in **Fig. 4C**, for all LEA contacts in V1, we compared MUA spike rates during the indicated early, middle, and late portions of the ITI period with the average MUA spike rate over the "background" period. UOA-stimulation conditions are shown schematically at the top left of each panel. Data are plotted as relative cortical depth versus average early (**Left**), middle (**Center**), or late (**Right**) ITI response normalized to the average background response (shaded regions: s.e.m.). *Asterisks* indicate significant differences in ITI and background response (t-test). Across all, but the whole array/7.8V condition, channels that responded to UOA stimulation (cf. **Fig. 2D-I and Fig.4C**) tended to show a significant decrease in MUA spiking, relative to the average background rate, particularly in the early portion of the ITI. Only few contacts showed significant changes in the later portion of the ITI and for all, but one, the changes were decreases in MUA spiking relative to background. The one exception to this pattern was seen on the responsive (deep-) mid-layer contacts during whole array/7.8V stimulation. On these contacts, there was a sharp increase in activity relative to baseline in the (early) middle period of the ITI. However, there was no significant difference between late ITI activity and background activity during this condition, indicating that by the end of the long ITI period spiking activity had returned to baseline. Importantly, these results indicate that the increases in MUA activity and their specific laminar patterns induced by UOA activation cannot be explained by thermal artifacts (see also **Extended Data Fig. 6**).
