## Supplemental Figure 6 for "An Optrode Array for Spatiotemporally Precise Large-Scale Optogenetic Stimulation of Deep Cortical Layers in Non-human Primates"

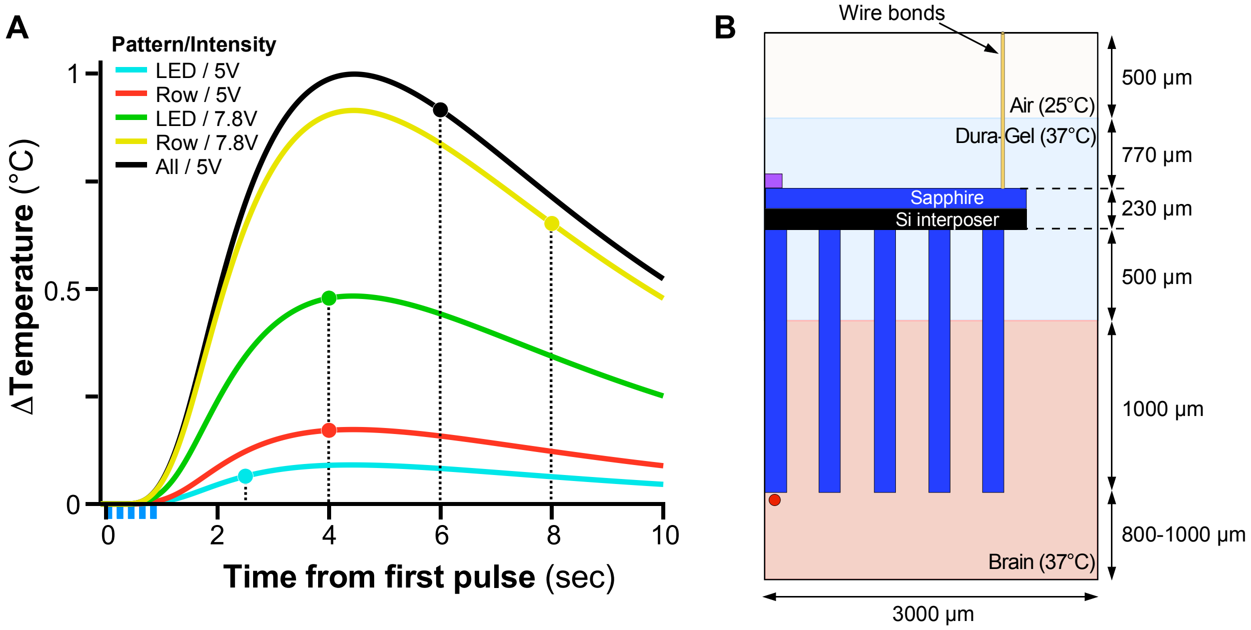


C

| Material | Thermal Conductivity (W/m/°C) | Heat Capacity (J/kg/°C) | Density (kg/m$^3$) |
|---|---|---|---|
| Brain | 0.55 | 3700 | 1040 |
| Dura-Gel | 0.2 | 840 | 970 |
| Silicon | 148 | 710 | 2330 |
| Sapphire | 40 | 700 | 3980 |
| Glass needle | 1 | 800 | 2500 |
| Gold (wire bonds) | 301 | 128 | 19320 |

**Extended Data Figure 6**

**Simulated temperature near the UOA tips (in L4C) under different photostimulation conditions.**
**(A)** Simulated temperature time profile in tissue under different operating conditions of the UOA, including a single µLED (*LED*), an entire row of µLEDs (*Row*), or the entire UOA (*All*), under different light intensities (5V or 7.8V). Heat generation occurs on the topside of the device and, due to the intervening Dura-Gel layer and tissue (superficial cortical layers 1-3), there is a delay in temperature rise at the stimulation site in L4 (*red dot* in panel B). This delay is approximately equal to the optical pulse stimulation period (five pulses at 100ms on and 100ms off, denoted as *blue bars* below the abscissa; the last optical pulse turns off at 0.9 sec). Temperature continues to rise during the first few seconds of the inter-trial interval (ITI) (the end of each ITI is marked by *colored dots;* longer ITIs were used for larger area and higher stimulation intensities), and peaks at ≤1°C for all conditions. Pulse pile-up is observed in certain conditions, resulting in a maximum temperature 31% ± 1% higher than the peak temperature observed after the initial pulse. **(B)** Illustration of the simulation geometry, showing the UOA glass needles, Si interposer layer, Sapphire substrate, and µLED (*purple*, only one illustrated). Gold wire bonds were also included (only one illustrated). Since the device was only partially inserted, the non-inserted portion is embedded in Dura-Gel. The air region was held at 25°C, the right-most Dura-Gel and brain boundaries were held at 37°C, and the lower brain boundary was held at 37°C (essential boundary conditions). A rotational symmetry boundary condition was used along the left edge. **(C)** Relevant properties of the constituent materials. A perfusion rate of 60 ml/min per 100g of tissue was included for brain.
