## Supplemental Figure 7 for "An Optrode Array for Spatiotemporally Precise Large-Scale Optogenetic Stimulation of Deep Cortical Layers in Non-human Primates"

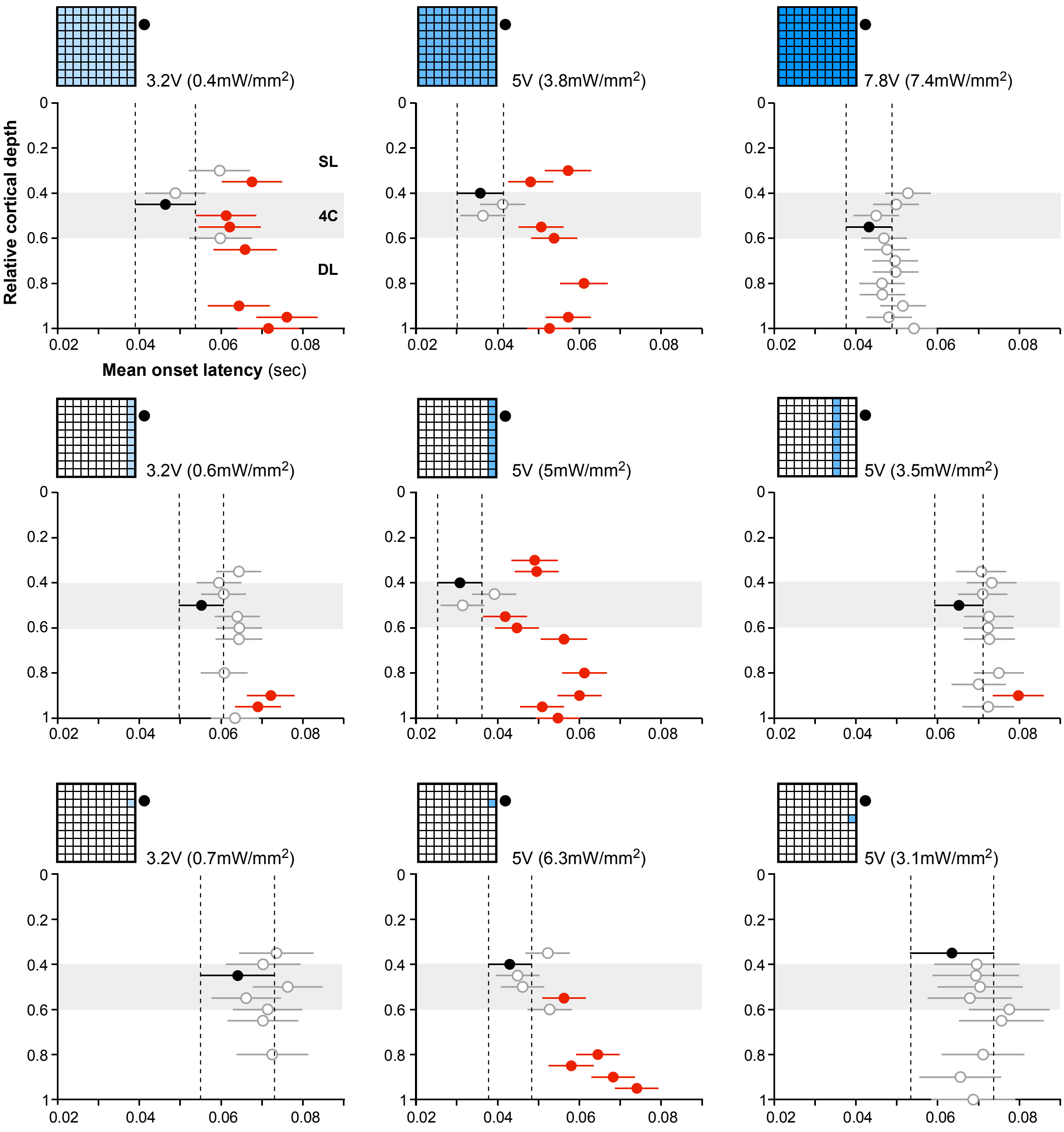


### Extended Data Figure 7

**Statistical analysis of onset latencies for penetration 2 (data shown in Figure 4C)**
Mean onset latency (± s.e.m) for each contact in P2 which showed significant response to UOA stimulation, for the UOA stimulation condition indicated by the *insets* at the top left of each plot. The mean latency was estimated from distributions of single-trial latency estimates. The *black dot* indicates the contact with the shortest latency in each condition. The *red dots* indicate the contacts that showed a statistically significant (Tukey HSD test) pairwise difference with the shortest latency contact (*black dot*), and the *empty dots* the contacts that did not differ significantly from the black dot. The *vertical dashed lines* indicate the points beyond which comparisons are significant.
