## Supplemental Table 1 for "An Optrode Array for Spatiotemporally Precise Large-Scale Optogenetic Stimulation of Deep Cortical Layers in Non-human Primates"

**Extended Data Table 1**

**Measured Mean Output Photostimulation Intensities for Different Input Voltages**

**WHOLE ARRAY**

| **Input Voltage**  (V) | **Output Optical Power**  (mW) | | | | | |
| --- | --- | --- | --- | --- | --- | --- |
|  | ***Mean*** | ***SD*** | ***Median*** | ***Min*** | ***Max*** | ***IQR*** |
| 2.8 | 0.0022 | 0.0016 | 0.0019 | 0 | 0.010 | 0.0014 |
| 3 | 0.0057 | 0.0040 | 0.0050 | 0.0005 | 0.024 | 0.0051 |
| 3.2 | 0.011 | 0.0072 | 0.010 | 0.0010 | 0.042 | 0.0091 |
| 3.5 | 0.022 | 0.013 | 0.020 | 0.0024 | 0.075 | 0.0170 |
| 4 | 0.044 | 0.026 | 0.041 | 0.0075 | 0.13 | 0.0313 |
| 5 | 0.10 | 0.056 | 0.088 | 0.018 | 0.27 | 0.0629 |
| 7.8 | 0.19 | 0.09 | 0.18 | 0.039 | 0.42 | 0.12 |

| **Input Voltage**  (V) | **Output Irradiance**  (mW/mm^2^) | | | | | |
| --- | --- | --- | --- | --- | --- | --- |
|  | ***Mean*** | ***SD*** | ***Median*** | ***Min*** | ***Max*** | ***IQR*** |
| 2.8 | 0.08 | 0.06 | 0.07 | 0.01 | 0.38 | 0.05 |
| 3 | 0.21 | 0.14 | 0.20 | 0.02 | 0.91 | 0.19 |
| 3.2 | 0.41 | 0.27 | 0.38 | 0.04 | 1.56 | 0.33 |
| 3.5 | 0.82 | 0.49 | 0.75 | 0.09 | 2.79 | 0.62 |
| 4 | 1.67 | 0.95 | 1.53 | 0.28 | 4.98 | 1.16 |
| 5 | 3.79 | 2.08 | 3.33 | 0.67 | 9.88 | 2.48 |
| 7.8 | 7.4 | 3.19 | 6.9 | 1.45 | 15.6 | 4.46 |

**COLUMN 1**

| **Input Voltage**  (V) | **Output Optical Power**  (mW) | | | | | |
| --- | --- | --- | --- | --- | --- | --- |
|  | ***Mean*** | ***SD*** | ***Median*** | ***Min*** | ***Max*** | ***IQR*** |
| 2.8 | 0.0030 | 0.0016 | 0.0024 | 0.0010 | 0.0068 | 0.0013 |
| 3 | 0.0079 | 0.0046 | 0.0071 | 0.0019 | 0.0181 | 0.0043 |
| 3.2 | 0.0156 | 0.0082 | 0.0146 | 0.0040 | 0.0321 | 0.0091 |
| 3.5 | 0.0305 | 0.0146 | 0.0301 | 0.0089 | 0.0570 | 0.0170 |
| 4 | 0.0611 | 0.0276 | 0.0627 | 0.0181 | 0.1016 | 0.0354 |
| 5 | 0.1350 | 0.0597 | 0.1324 | 0.0400 | 0.2424 | 0.0873 |
| 7.8 | 0.2548 | 0.0981 | 0.2719 | 0.1016 | 0.4168 | 0.1411 |

| **Input Voltage**  (V) | **Output Irradiance**  (mW/mm^2^) | | | | | |
| --- | --- | --- | --- | --- | --- | --- |
|  | ***Mean*** | ***SD*** | ***Median*** | ***Min*** | ***Max*** | ***IQR*** |
| 2.8 | 0.11 | 0.06 | 0.09 | 0.04 | 0.25 | 0.05 |
| 3 | 0.29 | 0.17 | 0.27 | 0.07 | 0.67 | 0.16 |
| 3.2 | 0.58 | 0.31 | 0.54 | 0.15 | 1.19 | 0.34 |
| 3.5 | 1.13 | 0.54 | 1.12 | 0.33 | 2.11 | 0.63 |
| 4 | 2.26 | 1.02 | 2.32 | 0.67 | 3.77 | 1.31 |
| 5 | 5.00 | 2.21 | 4.90 | 1.48 | 8.98 | 3.23 |
| 7.8 | 9.43 | 3.63 | 10.07 | 3.77 | 15.44 | 5.23 |

**COLUMN 3**

| **Input Voltage**  (V) | **Output Optical Power**  (mW) | | | | | |
| --- | --- | --- | --- | --- | --- | --- |
|  | ***Mean*** | ***SD*** | ***Median*** | ***Min*** | ***Max*** | ***IQR*** |
| 2.8 | 0.0020 | 0.0010 | 0.0022 | 0.0005 | 0.0032 | 0.0021 |
| 3 | 0.0055 | 0.0023 | 0.0058 | 0.0019 | 0.0089 | 0.0043 |
| 3.2 | 0.0106 | 0.0044 | 0.0110 | 0.0032 | 0.0159 | 0.0078 |
| 3.5 | 0.0211 | 0.0080 | 0.0216 | 0.0081 | 0.0321 | 0.0119 |
| 4 | 0.0421 | 0.0145 | 0.0432 | 0.0181 | 0.0657 | 0.0143 |
| 5 | 0.0933 | 0.0303 | 0.0932 | 0.0443 | 0.1460 | 0.0232 |
| 7.8 | 0.1836 | 0.0550 | 0.1820 | 0.0981 | 0.2871 | 0.0202 |

| **Input Voltage**  (V) | **Output Irradiance**  (mW/mm^2^) | | | | | |
| --- | --- | --- | --- | --- | --- | --- |
|  | ***Mean*** | ***SD*** | ***Median*** | ***Min*** | ***Max*** | ***IQR*** |
| 2.8 | 0.07 | 0.04 | 0.08 | 0.02 | 0.12 | 0.08 |
| 3 | 0.20 | 0.09 | 0.22 | 0.07 | 0.33 | 0.16 |
| 3.2 | 0.39 | 0.16 | 0.41 | 0.12 | 0.59 | 0.29 |
| 3.5 | 0.78 | 0.30 | 0.80 | 0.30 | 1.19 | 0.44 |
| 4 | 156 | 0.54 | 1.60 | 0.67 | 2.43 | 0.53 |
| 5 | 3.46 | 1.12 | 3.46 | 1.64 | 5.40 | 0.86 |
| 7.8 | 6.80 | 2.03 | 6.74 | 3.63 | 10.63 | 0.75 |

**COLUMN 5**

| **Input Voltage**  (V) | **Output Optical Power**  (mW) | | | | | |
| --- | --- | --- | --- | --- | --- | --- |
|  | ***Mean*** | ***SD*** | ***Median*** | ***Min*** | ***Max*** | ***IQR*** |
| 2.8 | 0.0014 | 0.0011 | 0.0010 | 0.0003 | 0.0041 | 0.0008 |
| 3 | 0.0041 | 0.0028 | 0.0032 | 0.0005 | 0.0097 | 0.0035 |
| 3.2 | 0.0081 | 0.0051 | 0.0068 | 0.0024 | 0.0181 | 0.0070 |
| 3.5 | 0.0165 | 0.0110 | 0.0142 | 0.0076 | 0.0360 | 0.0127 |
| 4 | 0.0344 | 0.0177 | 0.0249 | 0.0168 | 0.0713 | 0.0232 |
| 5 | 0.0797 | 0.0376 | 0.0683 | 0.0365 | 0.1583 | 0.0460 |
| 7.8 | 0.1623 | 0.0620 | 0.1456 | 0.0824 | 0.2971 | 0.0651 |

| **Input Voltage**  (V) | **Output Irradiance**  (mW/mm^2^) | | | | | |
| --- | --- | --- | --- | --- | --- | --- |
|  | ***Mean*** | ***SD*** | ***Median*** | ***Min*** | ***Max*** | ***IQR*** |
| 2.8 | 0.05 | 0.04 | 0.04 | 0.01 | 0.15 | 0.03 |
| 3 | 0.15 | 0.11 | 0.12 | 0.02 | 0.36 | 0.13 |
| 3.2 | 0.30 | 0.18 | 0.25 | 0.09 | 0.67 | 0.26 |
| 3.5 | 0.61 | 0.36 | 0.53 | 0.28 | 1.33 | 0.47 |
| 4 | 1.27 | 0.66 | 1.09 | 0.62 | 2.64 | 0.86 |
| 5 | 2.95 | 1.39 | 2.53 | 1.35 | 5.87 | 1.7 |
| 7.8 | 6.03 | 2.30 | 5.39 | 3.05 | 11.00 | 2.41 |

**SINGLE µLEDs IN COLUMN 1**

| **Input Voltage**  (V) | **Mean Output Optical Power**  (mW) | | | | | | | | |
| --- | --- | --- | --- | --- | --- | --- | --- | --- | --- |
|  | ***Row 1*** | ***Row 2*** | ***Row 3*** | ***Row 4*** | ***Row 5*** | ***Row 6*** | ***Row 7*** | ***Row 8*** | ***Row 9*** |
| 2.8 | 0.0024 | 0.0011 | 0.0024 | 0.0068 | 0.0019 | 0.0019 | 0.0032 | 0.0032 | 0.0046 |
| 3 | 0.0076 | 0.0019 | 0.0068 | 0.0181 | 0.0046 | 0.0041 | 0.0068 | 0.0089 | 0.0124 |
| 3.2 | 0.0146 | 0.0040 | 0.0146 | 0.0322 | 0.0089 | 0.0076 | 0.0138 | 0.0181 | 0.0246 |
| 3.5 | 0.0287 | 0.0089 | 0.0316 | 0.0570 | 0.0181 | 0.0159 | 0.0265 | 0.0351 | 0.0487 |
| 4 | 0.0562 | 0.0181 | 0.0691 | 0.1010 | 0.0365 | 0.0338 | 0.0521 | 0.0719 | 0.1016 |
| 5 | 0.1243 | 0.0400 | 0.1406 | 0.1938 | 0.0819 | 0.0824 | 0.1138 | 0.1697 | 0.2425 |
| 7.8 | 0.2016 | 0.1016 | 0.2857 | 0.3035 | 0.1597 | 0.1624 | 0.2581 | 0.2957 | 0.4168 |

| **Input Voltage**  (V) | **Mean Output Irradiance**  (mW/mm^2^) | | | | | | | | |
| --- | --- | --- | --- | --- | --- | --- | --- | --- | --- |
|  | ***Row 1*** | ***Row 2*** | ***Row 3*** | ***Row 4*** | ***Row 5*** | ***Row 6*** | ***Row 7*** | ***Row 8*** | ***Row 9*** |
| 2.8 | 0.1 | 0 | 0.1 | 0.3 | 0.1 | 0.1 | 0.1 | 0.1 | 0.2 |
| 3 | 0.3 | 0.1 | 0.3 | 0.7 | 0.2 | 0.2 | 0.3 | 0.3 | 0.5 |
| 3.2 | 0.5 | 0.2 | 0.5 | 1.2 | 0.3 | 0.3 | 0.5 | 0.7 | 0.9 |
| 3.5 | 1.1 | 0.3 | 1.2 | 2.1 | 0.7 | 0.6 | 1 | 1.3 | 1.8 |
| 4 | 2.1 | 0.7 | 2.6 | 3.7 | 1.4 | 1.3 | 1.9 | 2.7 | 3.8 |
| 5 | 4.6 | 1.5 | 5.2 | 7.2 | 3 | 3.1 | 4.2 | 6.3 | 9 |
| 7.8 | 7.5 | 3.8 | 10.6 | 11.2 | 5.9 | 6 | 9.6 | 11 | 15.4 |
